# Insights into CPSFL1 Induced Membrane Dynamics: A Multifaceted Regulator Linking Vesicle Formation to Thylakoid Biogenesis

**DOI:** 10.1101/2025.03.13.642960

**Authors:** Mastoureh Sedaghatmehr, Shreya Pramanik, Rumiana Dimova, Alexander Erban, Joachim Kopka, Alexander P. Hertle

**Affiliations:** Max-Planck Institute of Molecular Plant Physiology, Am Mühlenberg 1, 14476 Potsdam, Germany; Heinrich-Heine-Universität Düsseldorf, 40225 Düsseldorf, Germany; Max Planck Institute of Colloids and Interfaces, Science Park Golm, 14476 Potsdam, Germany; Cluster of Excellence on Plant Sciences, Heinrich Heine University, Düsseldorf, Germany; Oregon Health and Science University, Portland, Oregon, USA

## Abstract

Light drives plant life through photosynthesis, a process that takes place in the thylakoid membrane of the chloroplast, an organelle of cyanobacterial origin. The formation of thylakoid membranes within the chloroplast involves the eukaryote-specific factor *CHLOROPLAST SEC14 LIKE PROTEIN 1* (CPSFL1), which shares strong sequence homology with the vesicle trafficking regulator SEC14. CSPFL1 is essential for vesicle formation, yet its specific molecular function in this process has remained unclear. In this study, we characterized CSPFL1 functions both in vitro and in vivo. Using a minimal membrane system of giant unilamellar vesicles (GUVs), we show that CPSFL1 alone can induce vesiculation. This process is mediated by lipid binding and membrane deformation, driven by curvature sensing and lipid-protein electrostatics. When expressed in the prokaryote *E. coli*, the eukaryote-specific CSPFL1 induces membrane curvature and vesicle formation. Plastid CPSFL1 co-purifies with vesicular structures. Lipid compositional analysis of CPSFL1-induced vesicles from bacteria reveals the presence of quinone precursors as cargo, linking CSPFL-mediated vesicle formation to prenylquinone transport. Together, our data suggest that during plant evolution, the eukaryotic vesicle formation system was co-opted for the transport of membrane integral metabolites from the inner envelope to the thylakoid membrane.

## Introduction

Originating from a prokaryotic ancestor, plastids show remarkable versatility in both structure and function. The chloroplast is a specialized type of plastid in which photosynthesis occurs. The light reactions take place in the thylakoid membranes^1^. Unlike other membrane systems, thylakoid membranes predominantly consist of two galactolipids, monogalactosyldiacylglycerol (MGDG) and digalactosyldiacylglycerol (DGDG), as well as the sulfolipid sulfoquinovosyldiacylglycerol (SQDG), with relatively low phospholipid content ^2–4^. In addition, thylakoid membranes also contain other lipophilic substances, many of them derived from terpenoids. These include plastoquinone involved in electron transport and carotenoids involved in photoprotection ^5–6^.

Thylakoid membranes lack lipid synthesis activity entirely^5^. Most thylakoid lipids are synthesized at the chloroplast envelope from fatty acid species produced in the chloroplast stroma. Some lipids are imported across the envelope membranes from the ER^6^. Also, the key enzymes of quinone and carotenoid biosynthesis are exclusively located at the inner chloroplast envelope membrane^7,8^. How these components are then transported to the thylakoids is largely unknown.

The formation of thylakoid membranes from lipids, pigments, and proteins involves their synchronized synthesis in a precisely orchestrated spatiotemporal manner^9–12^. In many cases disruption in this intricate interplay results in photooxidative effects^13^. These results in damaged chloroplast membranes^14^. Also, deficiency of thylakoid membrane biosynthesis and the inability to assemble functional thylakoid membranes causes inability to establish photoautotrophism. Thus, many thylakoid biogenesis mutants are embryo or seedling lethal^15^. Only in a few cases, mutants can be partially rescued by growth on sucrose-containing media. These mutants are light sensitive, pale or white caused by either compromised thylakoid development or pleiotropic effects^9,12,16,23^. These phenotypes often hamper the functional characterization of respective factors *in vivo*. Therefore, synthetic and heterologous systems serve as useful alternatives ^31–36^.

The chloroplast localized Sec14-like protein 1 (CPSFL1), is also essential for plant development and plays a key role in thylakoid biogenesis ^16–18^. *Arabidopsis* plants lacking CSPFL1 are seedling lethal but survive, when grown heterotrophically^32^. Mutant chloroplasts show reduced amounts of thylakoid membranes and these exhibit a disordered simplified organisation with less prominent grana stacking and reduced number of interconnected stroma lamellae as compared to wild type (WT) thylakoids^16,18^. Vesicle transport, contact sites, and lipid transport proteins were proposed as mechanisms for shuttling lipids and terpenoid derivates from the envelope to the thylakoid bilayer^19–22^. In *cpsfl1-1*, chloroplast stromal vesicles cannot be detected^16,23^. Instead, remaining thylakoids are directly connected to envelope membranes. Since contact sites between thylakoid and envelope membranes are rare in mature chloroplasts of WT plants, the increased occurrence of contact sites in *cpsfl1* mutants suggests that there is a defect in thylakoid separation^16^. Nevertheless, these direct connections likely also serve to supply the residual thylakoids directly with lipids and isoprenoid components. However, the remaining thylakoids of *cpsfl1* mutants exhibit a significantly lower proportion of carotenoids and quinones^17,18^. There are multiple conceivable explanations for this observation. Either mutants are directly affected in terpenoid biosynthesis^18^ or they have a defect in a mechanism that transports these substances from their site of synthesis (i.e. the inner chloroplast envelope) to the thylakoid membrane. The former is supported by the decreased expression of enzymes involved in tetrapyrrole, quinone and carotenoid biosynthesis^18^. However, changes in nuclear gene expression could be a secondary effect of the strong phenotype.

The chloroplast localized Sec14-like protein 1 (CPSFL1) impacts thylakoid biogenesis likely through both, vesicle transport and isoprenoid metabolism ^16–18^. The mechanistic framework remained so far unclear. The name-giving Sec14 protein from yeast, regulates vesicle transport between the *trans*-Golgi network and the plasma membrane by influencing the levels of polyphosphatidylinositides (PPIs)^24^. CPSFL1 can complement the yeast sec14 mutation^16^. *In vitro*, CPSFL1 binds directly to PPIs and phosphatidic acid (PA) and mediates PPI transport directly as a lipid transfer protein (LTP)^16,18^.

Both vesicle transport and transfer of lipids across contact sites involves membrane deformation and the generation of curvature^25^. For yeast Sec14, a stronger curvature of the membrane promotes both membrane binding and lipid exchange properties^26^. A common principle of membrane bending by proteins is the insertion of an amphiphilic helix (AH) into the membrane^27–29^. Proteins like endophilins, amphiphysins, epsins and CURT1 cause initial membrane curvature by insertion of their amphiphilic helix into only one-half bilayer leaflet^29–32^. This imposes curvature to the membrane by increasing the area of one leaflet over the other^29^. Scaffold forming proteins, like clathrins or VIPP1, oligomerize to form a rigid structure^33–35^. Rigid multiprotein assemblies bind and deform the underlying membrane^33^. Lipids also play a crucial role in inducing membrane curvature^36–38^. They serve as signal to recruit membrane deforming proteins^39^. In addition, locally leaflet specific accumulation of cone-shaped lipids (i.e. PA, DAG or PE) or inverted cone-shaped lipids (i.e. PPIs and Lyso-PA) can deform membranes via their conical shape^40,41^. Such increase can be achieved by upregulation of lipid synthesis or oligomerisation of proteins that bind these specific lipids and thus enrich them locally. In bio-membranes, a balanced mixture of bilayer forming lipids and non-bilayer lipids, is actively maintained^42,43^. This allows them to form stable bilayers most of the time but also to be subject to disruption during localized events such as membrane fusion, endocytosis and cell fission^44^. While bilayer lipids spontaneously arrange into flat bilayers, non-bilayer lipids such as PE in bacterial and MGDG in plastid membranes do not easily form bilayers and instead tend to adopt other types of structures, such as hexagonal or cubic phases. Such localized events can be triggered by protein-lipid interactions and cause local changes in electrostatics and hydration which result in L_α_ to H_II_ phase transition of cone-shaped lipids. This in turn affects membrane curvature and locally destabilizes the lipid bilayer^45^. These events may trigger repair mechanisms or stress responses potentially involving ESCRTIII and SEC14 proteins.

In order to understand a complex multifactorial process such as thylakoid biogenesis, the use of biomimetic systems has proven useful. These allows to study isolated aspects of membrane protein interaction, protein trafficking and lipid membrane formation without pleiotropic effects Biomimetic systems in the form of giant unilamellar vesicles (GUVs)^46^ have proved extremely useful for investigating shape deformation, budding and division of membranes^45,47–49^. So far thylakoid-mimicking GUVs have not yet been reported.

Here we construct minimalistic GUV-based models of thylakoid-like membranes and show, that CPSFL1 interacts with synthetic and bio membranes consisting of diverse lipid stoichiometries based on a curvature sensing mechanism. CPSFL1 forms oligomers to deform and transport large amounts of membranes in vesicles. While the heterologous system in *E. coli* demonstrates transport of metabolites dissolved in the membrane or enclosed in vesicles the nature of the cargo in plant chloroplasts remains to be explored.

## Results

### Minimal GUV models of thylakoid membranes uncover CPSFL1 dependence on its amphiphilic helix and anionic lipids for binding

CPSFL1 transports lipids and is found in chloroplast fractionations mainly as a soluble protein^16^. However, when overexpressed, it has also been localized to chloroplast membranes^18^. To understand the structural features of CPSFL1’s membrane association, we analysed the interaction of recombinant CPSFL1 with synthetic membranes (Fig. 1). Comparative *in silico* analyses with its yeast homologue Sec14 using AlphaFold, together with helical wheel predictions using HeliQuest suggested that CPSFL1 forms an amphiphilic helix (Fig. 1a, supplemental Figure 1a). Such helices facilitate protein membrane interactions.

**Figure 1:**
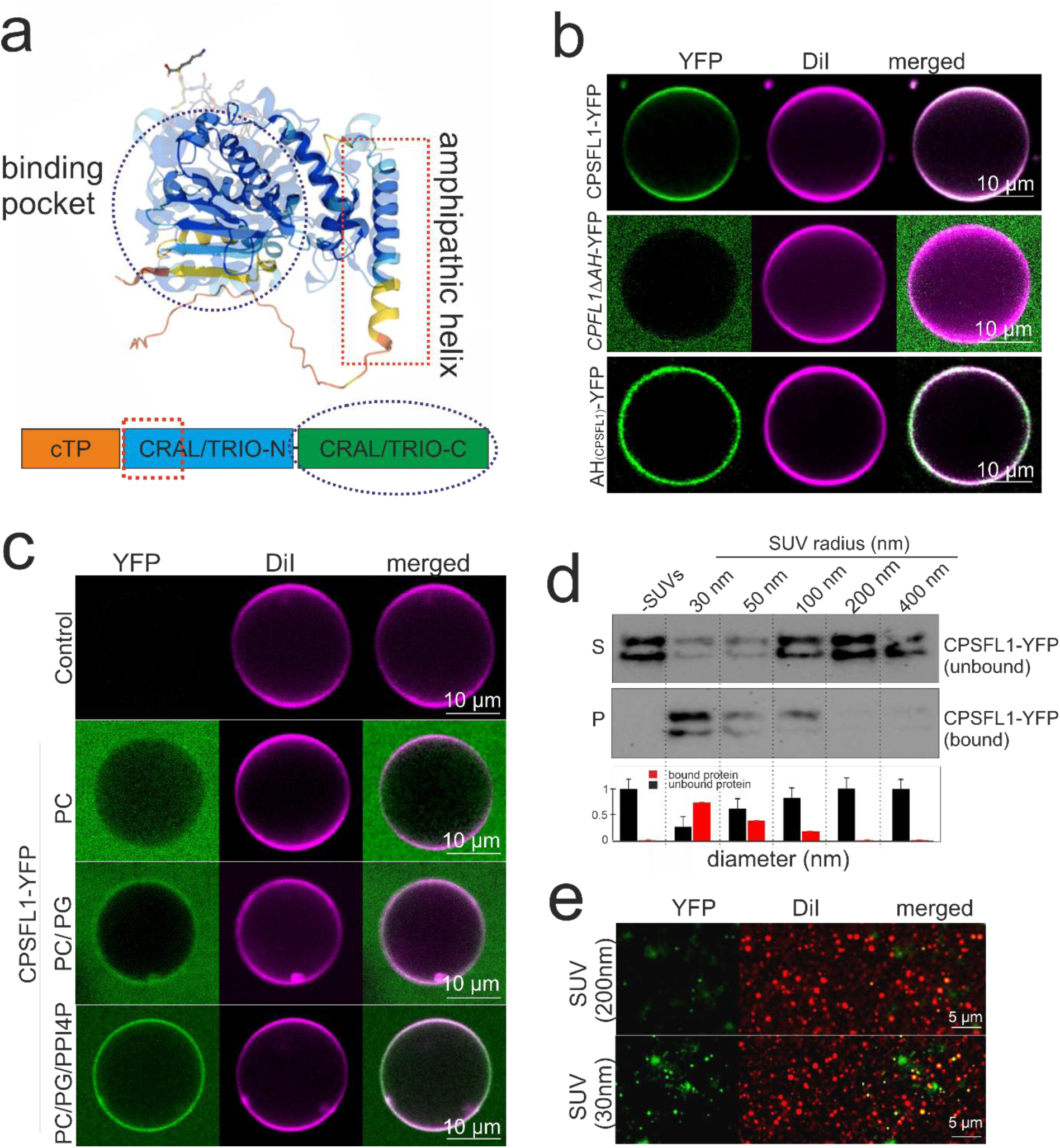
Membrane binding, curvature sensing and structural analysis of CPSFL1. **a,** Comparison of yeast Sec14 and CPSFL1 using superimposed structures predicted by AlphaFold. Regions in blue represent most confident predictions, whereas yellow to orange are regions of low confidence. A prominent long α-helix highlighted within the CRAL TRIO_N domain (CRAL/TRIO_N) of CPSFL1 (red dotted line, amphipathic helix) following the chloroplast targeting peptide (cTP) apart from its lipophilic binding pocket within the CRAL TRIO_C domain (blue dotted line, binding pocket) is depicted in the scheme below. **b,** Membrane binding of recombinant YFP tagged CPSFL1 and mutant variants were analysed *in vitro* using GUVs with chloroplast specific lipid composition obtained via PVA assisted hydrogel swelling. Following addition of YFP tagged recombinant CPSFL1 protein variants (CPSFL1-YFP, *CPSFL1-ΔAH-YFP*, AH_(CPSFL1)_-YFP) to GUV suspensions (1 µM final protein concentration), protein fluorescence (YFP) and GUV fluorescence (DiI) was imaged and overlayed (merged) using confocal microscopy. A role of CPSFL1 amphipathic helix in membrane binding was confirmed. **c,** Recombinant fluorescent CPSFL1-YFP (1µM final) was added to GUVs formed from electroneutral phospholipid DOPC (5 mM lipid concentration), negatively charged GUVs (DOPC/DOPG mix 1:1) or GUVs with conical curvature inducing phospholipid (DOPC/DOPG/PI4P (50/49.9/0.1). Protein fluorescence (YFP) and GUV fluorescence (DiI) was imaged and overlayed (merged) using confocal microscopy. Moderate binding of CPSFL1 to charged lipids and strong binding to charged and conical lipid containing membranes was observed. GUVs with neutral surface were not bound by CPSFL1. **d**, Large (LUVs) and small unilamellar vesicles (SUVs) with decreasing diameters (200-30 nm) were prepared from phosphatidylcholine (PC)/phosphatidylglycerol (PG) /1,1’-Dioctadecyl-3,3,3’,3’-Tetramethylindocarbocyaninperchlorat (DiI) mixtures (1:1:0.01). Following co-incubation with CPSFL1-YFP protein (1 µM final) samples were fractionated by centrifugation and the amount of CPSFL1-YFP in supernatant (S) and pellet (P) fractions was analysed by SDS-PAGE and western blotting using GFP specific antibodies and quantified using ImageJ (lower panel). As compared to LUVs of 200 nm diameter, smaller SUVs (30 to 50 nm) co-purified much higher levels of CPSFL1-YFP. **e**, Pellet fractions of CPSFL1-YFP and LUVs or SUVs described in e, were analysed by fluorescence microscopy. Higher abundance of CPSFL1-YFP (YFP, protein) was found using SUVs with smaller diameters (30 nm) (DiI, lipid).

We first designed minimalistic GUV-based models of thylakoid-like membranes. The GUVs were composed of the key chloroplast thylakoid lipids MGDG, DGDG, PG, SQDG and PI in a molar ration of 52:26:6.5:9.5:1 ^2,4^. The vesicles were fluorescently labelled by adding 0.1 mol% DiI lipid dye into the lipid mixtures. Co-Incubation with recombinant CPSFL1-YFP showed the YFP signal colocalizing with the GUV fluorescence (Fig. 1b, CPSFL1-YFP, supplementary Fig.1b). This confirmed membrane recognition and binding of CPSFL1. The role of the amphiphilic helix in membrane binding was confirmed by expressing a deletion mutant of the amphiphilic helix (CPSFL1*ΔAH-YFP*) and the α-helix of CPSFL1 in isolation (*AH-_CPSFL1_-YFP*) (Fig.1b, supplementary Fig.1b). While the YFP signal associated with the GUV was strongly decreased in the former, the helix alone conferred a strong YFP signal colocalizing with the membrane (Fig. 1b, lower and middle panel, supplementary Fig.1b).

Chloroplast membranes are mainly composed of electroneutral galactolipids but also contain lower amounts of charged sulfo- and phospholipids, namely SQDG and PG^2–4^. Thus, we tested whether CPSFL1s membrane binding depends on electrostatic interactions by using charged phospholipids (Fig. 1c, supplementary Fig.1c). Liposomes composed of neutral lipids, such as DGDG or MGDG/DGDG, have been shown to aggregate in the absence of charged lipids^50^. Since both are electroneutral, we instead used PC as a neutral bilayer-forming lipid to assess initial binding in the absence of a net surface charge (Fig. 1c, control, supplementary Fig.1c). GUVs composed solely of PC did not interact with recombinant CPSFL1-YFP (Fig. 1c, PC, supplementary Fig.1c). Instead CPSFL1 remained in the soluble fraction, as indicated by the diffuse protein fluorescence surrounding the GUV (Fig. 1c, PC, YFP, supplementary Fig.1c). Supplementing the GUVs with anionic lipid PG in the bilayer mix introduced a net negative surface charge (PC/PG) (Fig. 1c, PC/PG, supplementary Fig.1c). These GUVs exhibited weak binding of CPSFL1 as indicated by both membrane-associated and surrounding YFP fluorescence (Fig. 1c, PC/PG, YFP supplementary Fig.1c). This suggests that electrostatic interactions contribute to membrane binding. However, unlike thylakoid-mimicking GUVs, a significant fraction of CPSFL1 remained unbound (Fig. 1b, CPSFL1-YFP and Fig. 1c. PC/PG YFP, supplementary Fig.1b, c), indicating that electrostatic interaction alone may not fully account for CPSFL1’s membrane binding affinity.

### CPSFL1 senses membrane curvature

Human and yeast Sec14 protein can recognize membranes through membrane curvature sensing^26,51^. To investigate whether CPSFL1 exhibits similar property, we tested its ability to sense membrane curvature (Fig. 1c, d and e). First, we induced packing defects in GUVs with diameters of around 50 µm (essentially exhibiting a flat membrane) by incorporating negative cone-shaped lipids into bilayer mixtures (Fig. 1c, supplementary Fig.1c). A small amount of PI4P was added to the lipid mix (DOPC/PI4P, 99.8:0.2) to introduce packing defects in GUV membranes. Upon addition of CPSFL1, strong membrane binding was observed (Fig. 1c, PC/PG/PI4P, supplementary Fig.1c). CPSFL1 exhibited a higher binding affinity to PI4P-containing GUVs compared to those containing only PG (Fig. 1c, PC/PG vs. PC/PG/PI4P, supplementary Fig.1c). Since PI4P is both, negatively charged and conical, these results suggest that CPSFL1 preferentially binds to less well-packed membranes.

To further test curvature sensing ability of CPSFL1, we employed an alternative approach using negatively charged large and small unilamellar vesicles (LUVs and SUVs) with varying curvatures (LUVs: 100-1000 nm in diameter; SUVs: <100 nm diameter) (Fig. 1d, e). Since curvature is inversely proportional to vesicle radius, smaller vesicles (30 nm) exhibit higher curvature than larger ones (200 nm), while maintaining a constant lipid composition^48,52–54^. LUVs and SUVs were prepared from DOPC/DOPG (1:1) lipid mixtures and co-incubated with equal amounts of recombinant CPSFL1-YFP (1 µM final concentration). Following liposome sedimentation, we assessed CPSFL1 co-sedimentation both immunologically and via confocal microscopy (Fig. 1d, e). Immunologic analysis showed an increased co-sedimentation of CPSFL1 with decreasing liposome diameters (Fig.1 d). Microscopic examination of re-isolated SUVs co-incubated prior with CPSFL1, further confirmed significantly higher levels of CPSFL1-YFP associated with smaller SUVs (30 nm) compared to larger LUVs (200 nm) (Fig. 1e). All together, these results indicate that CPSFL1 recognizes membranes through a combination of electrostatic interactions and curvature-dependent membrane binding.

### CPSFL1 induces GUV deformations and vesicle budding

Oligomerisation of membrane associated or lipid binding proteins on membranes often results in lipid sorting, bilayer deformation or remodeling^52,53^. As most dramatic consequence, membranes rupture or form nanopores^54^. Thus, we investigated the impact of recombinant CPSFL1-YFP on the appearance of GUVs with chloroplast inner envelope lipid compositions over time by using confocal microscopy (Fig. 2). Initially GUVs co-incubated with CPSFL1 remained spherical for several minutes (Fig. 2 a, initial). However, a continuous and moderate deformation of the GUVs in the form of inward invagination (partial budding) was observed following longer co-incubation times (Fig. 2 a, +10 min). This was visible by both, DiI membrane and CPSFL1-YFP protein fluorescence (Fig. 2 DiI, YFP). Furthermore, deformation was asymmetric and characterized by negative curvature (the induced dents were curved inward) and accompanied by volume loss. In contrast, raising external osmolarity of the GUVs in the absence of CPSFL1 lead to volume loss and smooth deflation without preferred direction of the curvature (Fig.2b, +deflation). No fluctuation of membrane shape indicated a rather rigid membrane of the vesicles. Thus, the inward dents observed on the GUVs indicate that CPSFL1 actively induces asymmetric membrane deformation. No lipid phase separation or complete budding of vesicles was observed under these conditions. Instead, fluorescence microscopy revealed homogeneous distribution of both proteins and lipids (Fig. 2a, YFP). However, over time the overall background fluorescence in protein and lipid channels increased (Fig. 2c, background, supplementary Fig. 2a, green graph). Simultaneously, DiI mediated membrane fluorescence decreased indicating membrane loss or photobleaching effects (Fig. 2c, DiI).

**Figure 2:**
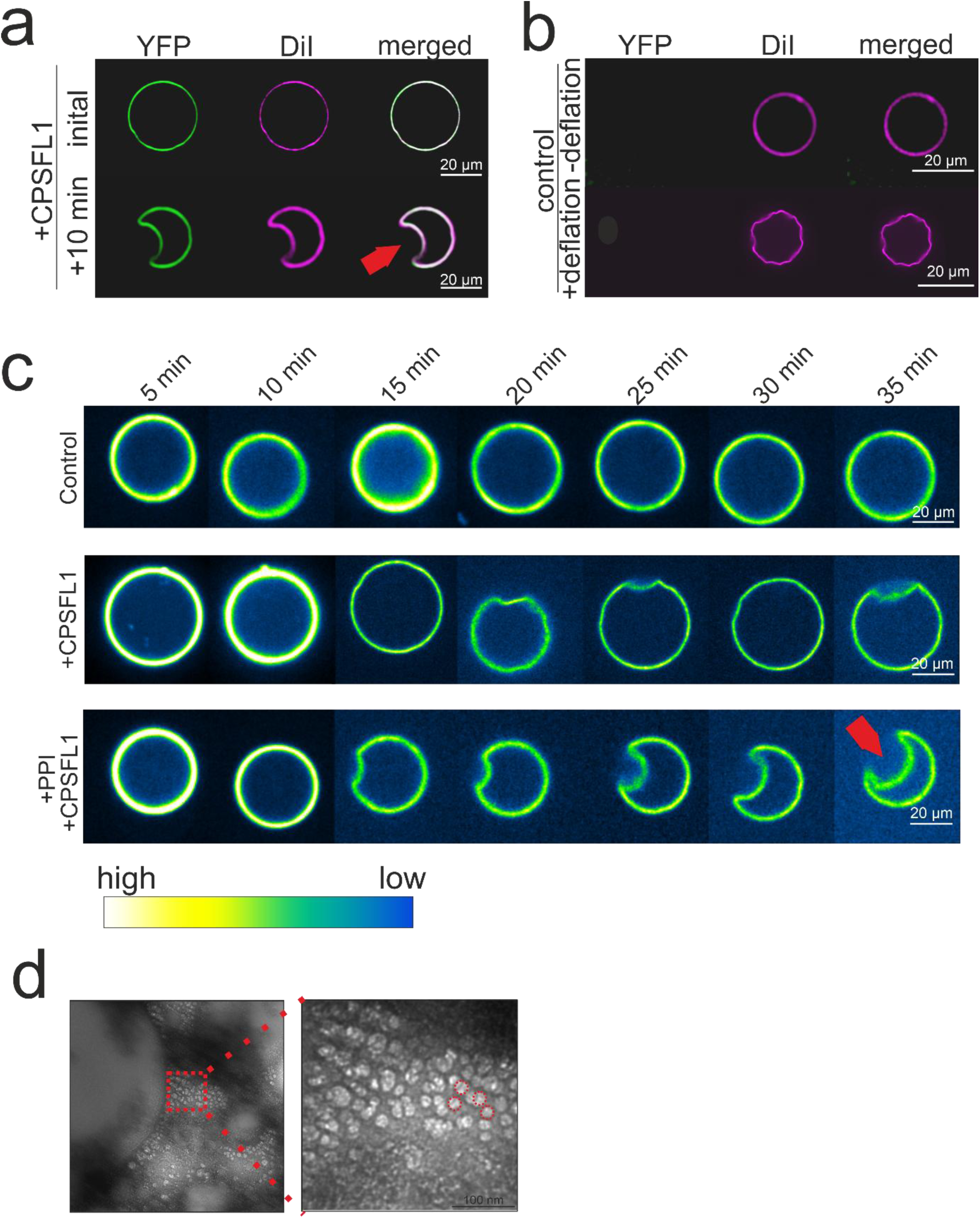
Membrane curvature modulation and vesicle formation by CPSFL1. **a,** GUVs made from MGDG, DGDG, PG, SQDG and PI in a molar ration of 52:26:6.5:9.5:1stained with DiI for fluorescence detection were co-incubated with recombinant CPSFL1-YFP proteins to study membrane binding. Co-incubation was analysed by confocal microscopy detecting recombinant CPSFL1-YFP (YFP fluorescence, green) and GUVs (DiI fluorescence, magenta) as co-localized. Observation of the same spherical GUV with bound CPSFL1-YFP (initial) showed a moderate deformation after several minutes (+10 min.). **b,** As a control GUVs made from synthetic chloroplast lipids (control) were stained by DiI (magenta) and analysed by confocal microscopy without CPSFL1. GUVs remained stable and without phase separation. Increasing the osmolarity of the GUVs containing buffer via evaporation lead to a volume loss and symmetric deflation of GUVs. **c,** Changes of fluorescence intensity (intensity range indicator) and shape of GUVs was observed over time (each picture represents 5 min time interval). As visualized by false-coloured images of GUV fluorescence with range indicator. Control experiments using heat denatured recombinant protein solutions showed no significant differences over time (control, lower panel). Left: Quantification of lipid fluorescence over time. Following CPSFL1-YFP addition, GUV deformation was accompanied by decrease in the fluorescence of the membranes (magenta) and an increase in the fluorescence of the background (green). In control experiments fluorescence intensities remained unchanged. **d,** GUVs were analysed in higher resolution following negative staining by TEM analysis on CPSFL1 treated GUVs. Numerous spherical structures were observed next to GUVs indicating CPSFL1 aggregation or vesicle budding.

To rule out photobleaching or contamination with co-purified components (e.g., detergents, lipophilic molecules), control experiments were conducted using identical but heat-inactivated CPSFL1 fractions (Fig. 2c, Control, supplementary Fig. 2a, magenta line). No membrane deformation was observed in these controls (Fig. 2b and c, control) and DiI fluorescence remained unchanged (Fig. 2b, supplementary Fig. 2a, magenta graph), confirming that the observed effects were not due to bleaching.

Next, we examined the influence of the conical lipid PI4P on CPSFL1-dependent membrane deformation (Fig. 2c, +PPI, supplementary Fig.2b). Following addition of CPSFL1 caused an even more pronounced inward bending of the GUVs resembling a dumbbell shape (Fig. 2c, +PPI+CPSFL1). Again, DiI fluorescence in the GUV surrounding soluble fraction increased while membrane fluorescence decreased (Fig. 2 d, right, supplementary Fig.2b, green graph). In contrast, fluorescence in control experiments remained unaltered (Fig.2 d, supplementary Fig.2b, magenta graph). These CPSFL1-dependent fluorescence changes in the solution around the GUVs suggest an active mechanism of membrane loss.

To further investigate this process at higher resolution, we analysed CPSFL1-treated GUVs using negative staining and TEM (Fig. 2d). Surprisingly, numerous small vesicles surrounding the GUVs were observed, raising the possibility that CPSFL1 mediates vesicle or membrane particle formation when applied externally (Fig. 2d, left). Next, we investigated the effect of CPSFL1 expression within a living system.

### Heterologous expression of CPSFL1 induced an intracellular lipophilic compartment composed of numerous vesicles

To explore the consequences of CPSFL1 expression in a “living” bacterial context, we used *E. coli* as a model organism (Fig. 3). The inducible expression of CPSFL1 in *E. coli* did not significantly affect growth rate or cell shape. Still, analysing cryo-fixed CPSFL1 expressing cells with electron microscopy revealed ultrastructural changes (Fig. 3a). We observed internal vesicles, cell membrane deformation and a distinctive intracellular osmiophilic compartment indicating a lipophilic environment (Fig. 3a, red arrows). Importantly, these effects were not observed when a control protein (C-terminus of KEA3^55^) with similar properties were expressed in *E. coli* (supplementary Fig. 3a). These results underscore CPSFL1’s specific impact on cellular architecture and its property in influencing membrane dynamics *in vivo*. To verify whether CPSFL1 is directly associated with the observed structural changes, we determined its subcellular distribution by immunolocalization using FLAG specific antibodies (Fig. 3b). This revealed CPSFL1’s localisation as membrane-associated and soluble protein. Furthermore, CPSFL1 localised also at the outer edges and within the dark osmiophilic structures (Fig. 3b, lower left). Additionally, gold particle clusters in the soluble fraction hinted oligomerisation into multidomain protein polymers (Fig. 3b, lower images). Altogether, this suggests a membrane associated and soluble fraction of CPSFL1 with highest abundance in a central osmiophilic compartment. To understand these structures in more detail, we purified CPSFL1-FLAG omitting any detergents and ions (e.g. Mg^2+^ or Ca^2+^) from *E. coli* lysates under native conditions. These protein preparations were subsequently analysed by negative staining and TEM (Fig. 3c). Here, larger lipophilic structures with a bramble-like surface were observed within the native protein fractions (Fig. 3c, top left). Closer examination resolved these and surrounding structures as composed of numerous small globular structrures (Fig. 3c, top right). These had diameters with an average of 3.57 nm (Fig. 3c, lower panel). Next, we conducted an in-depth compositional analysis of native CPSFL1 purifications by SDS-PAGE and colloidal Coomassie staining. CPSFL1 was identified by far as the main protein within these extracts (Fig. 3d, Coomassie, left lane). In addition, the molecular composition the lipophilic components co-purified with CPSFL1 was analysed using thin-layer chromatography (TLC) (Fig. 3d, right lane). The analysis predominantly identified a mixture of membrane lipids. As. As compared to total lipid extracts of *E. coli*, CPSFL1 purified fractions, had significantly decreased phosphatidyl ethanol (PE) content. Considering the majority of PE present in the outer bacterial membranes, our TLC analysis comparing total lipid extracts and recombinant CPSFL1 lipid extracts reveal PG, CL and PE stoichiometries typical of the inner bacterial membrane^55^ (Fig. 3e). This implies the possibility that the globular structures bound by CPSFL1 represent vesicles and stem from the inner membrane. This indicates a dynamic role for CPSFL1 in generating vesicular structures from the bacterial inner membrane.

**Figure 3:**
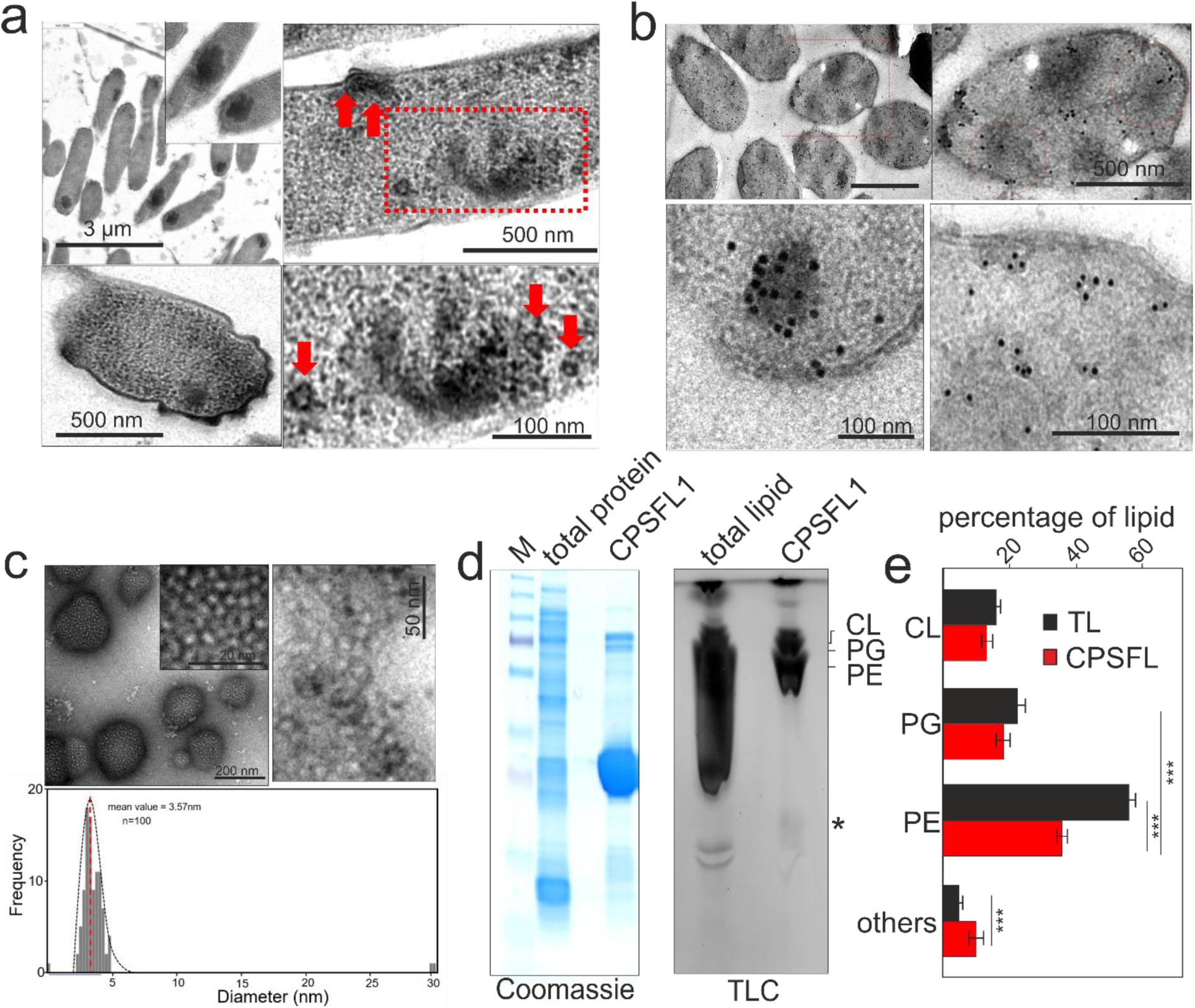
Expression of CPSFL1 in *E. coli* leads to ultrastructural changes and vesicle formation. **a,** Upon expression of CPSFL1-Flag in *E. coli* cells, a dark (osmiophilic) compartment appeared within the cells (red star indicates position of magnified area of inset, top left). Furthermore, deformations of the envelope membranes and vesicular structures within the cytoplasm were observed exclusively in CPSFL1 expressing cells (top and bottom left, red rectangle and arrows). **b,** Localization of CPSFL1-Flag in *E. coli* cells. Immunolabeling was performed using Flag specific antibodies. Gold particles were detected exclusively within the cells and at the membrane and in the soluble compartment (top). Particle clusters appeared within the osmiophilic area and within the cytoplasm indicating aggregation (bottom). Higher magnification of immunogold particle clusters in CPSFL1 expressing *E. coli* cells (bottom). **c**, Following purification of CPSFL1 under native conditions from *E. coli* cells, fractions were negatively stained and analysed by TEM. Prolamellar body like structures with a diameter of 200-500 nm were detected (upper panel and inset). Higher magnification identified these structures as vesicle assemblies (lower panel). Measurement of the vesicles identified an average diameter of 3,5 nm (lower). **d,** Compositional analysis of total and native purified CPSFL1-Flag protein and lipid fractions from *E. coli* cells was done by SDS-PAGE and TLC respectively. Proteins were visualized by colloidal Coomassie staining against marker proteins (M). Lipids were visualized by Cu^2+^-sulphate charring. (*) marks substance only found in CPSFL1 fraction. **e,** Quantification of lipid mixtures obtained following TLC, Cu^2+^-sulphate charring from total lipid extracts of *E. coli* (TL) and recombinant purified CPSFL1 fractions (CPSFL) shows a strong decrease in PE. PE: phosphatidylethanol, PG: phosphatidylglycerol, CL: cardiolipin.

### Probing membrane dynamics in *E. coli*: from vesicle formation to endocytosis

Previous experiments demonstrated CPSFL1 affinity to bind to small vesicles due to high curvature (Fig. 1d). Whether, CPSFL1 co-purified vesicles represent microsomal fractions formed by membrane fragmentation or represent actively formed vesicles was unclear (Fig. 3c). To confirm the nature of the observed vesicles and compartment, we stained membranes of *E. coli* cells with membrane-impermeable dye FM4-64. Internalization of FM4-64 upon CPSFL1 expression would indicate endocytosis (Fig. 4a, FM4-64). Alternatively, and due to the osmiophilic nature of the CPSFL1 induced compartment observed by TEM, we also used membrane-permissive lipophilic dye BODIPY (Fig. 4a, BODIPY). In contrast to FM4-64, this allows staining of lipophilic compounds in addition to membrane lipids as observed for lipid droplets or lipid phases (Fig. 4a, BODIPY). Upon protein production BODIPY labelled CPSFL1-induced subcellular structures indicated lipophilic and osmiophilic properties (Fig. 4a, green signal). Simultaneously, increased FM4-64 internalisation and staining of intracellular compartments was observed indicating endocytosis-like membrane transport (Fig. 4a, red signal). Yet, the fluorescent dyes did not show a complete overlay (Fig. 4a, merged image).

**Figure 4:**
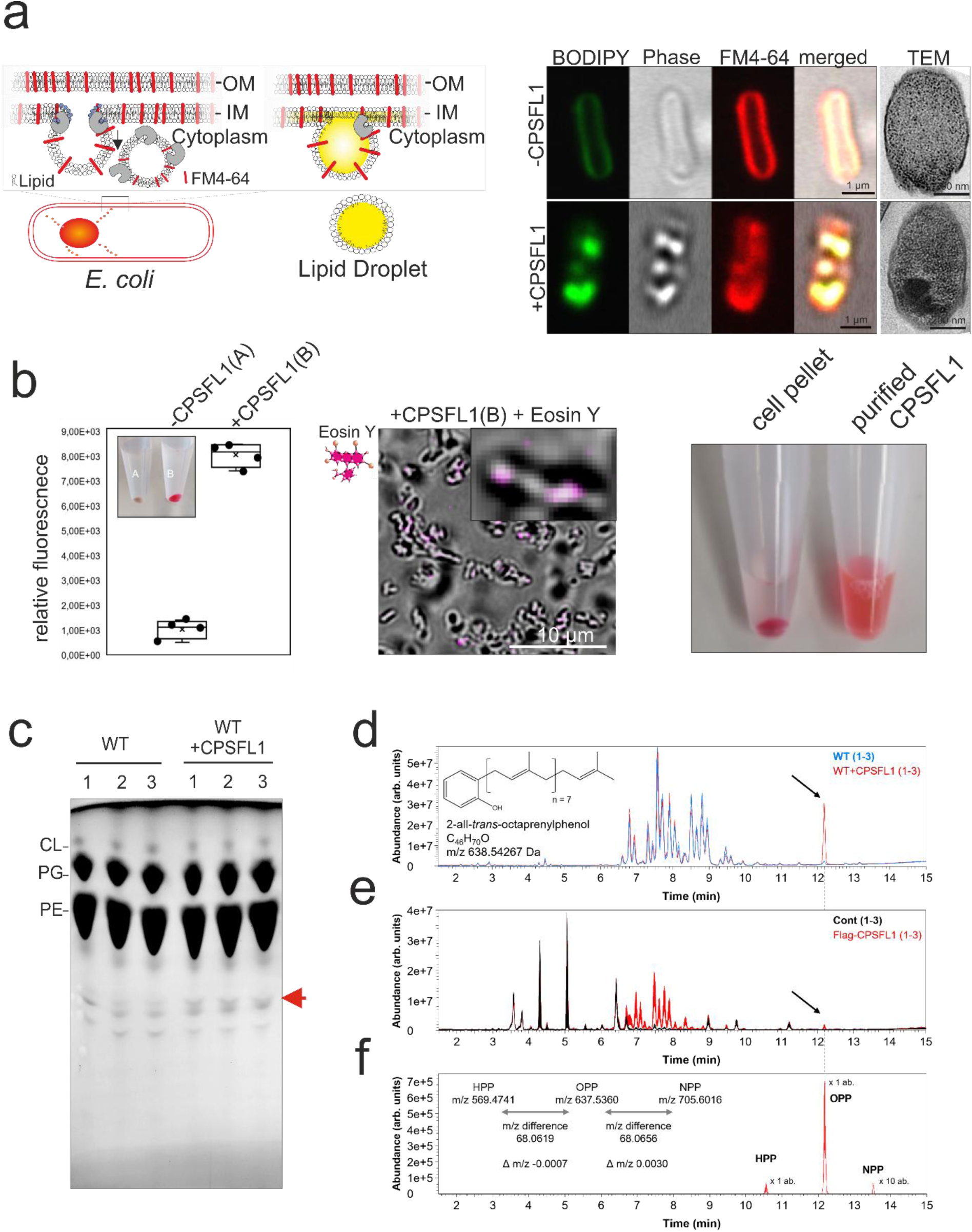
CPSFL1 induces endocytosis like vesicle formation. **a,** Schematic drawing of putative dye uptake into *E. coli* cells upon CPSFL1 expression (left). BODIPY dye uptake indicates formations of lipid droplets, FM-4-64 uptake indicates endocytosis of vesicles. Right, Uptake by endocytosis of dyes FM4-64 or BODIPY by *E. coli* cells expressing CPSFL1. Upon induction of CPSFL1 by IPTG FM4-64 located in envelope membranes becomes internalized. The lipophilic compartment is highly stainable with membrane permeable lipid dye BODIPY indicating the presence of neutral lipids or lipophilic compounds and also stains with osmium indicating presence of unsaturated fatty acids. Scale bar:500 nm. **b,** Uptake of fluorescent membrane impermeable water-soluble dye Eosin Y by *E. coli* cells from the periplasmic space following CPSFL1 expression. Left, dye was removed from the external medium by repeated washing and sedimentation. In control strains (-CPSFL1 (A)) washing removed the dye entirely (white pellet). However, cells expressing CPSFL1 retained the dye indicating binding or internalisation of fluorescent labelled dye by bacteria (+CPSFL1(B)) (pink pellet). Middle, Confocal image showing bacteria expressing CPSFL1 after washing in brightfield and Eosin Y dye fluorescence inside the cells. Right, *E. coli* cells following Eosin Y application and extensive washing remained pink. Eosin co-purifies with *E. coli* cells and co-purified with CPSFL1 following native purification. **c,** Comparative compositional analysis of lipophilic extracts obtained from WT and CPSFL1 expressing *E. coli* cells using TLC. Red arrow marks additional band appearing only when CPSFL1 was expressed. PE: phosphatidylethanol, PG: phosphatidylglycerol, CL: cardiolipin. **d,** Lipids of *E. coli* cells expressing flag-tagged CPSFL1 compared to control protein expressing cells. The differential display of 3 technical control LC-MS analyses (blue, overlay) and 3 analyses of a lipid preparation from flag-tagged CPSFL1 expressing cells (red, underlay) identifies accumulation of an abundant lipid (arrow). This substance was annotated as ubiquinone biosynthesis intermediate, octaprenylphenol (OPP, inserted structure represents *E. coli* metabolite, 2-all-*trans*-octaprenylphenol). Annotation was supported by exact mass, and co-accumulation of minor lipids matching by relative chromatographic retention and exact masses to hepta- and nonaprenyl phenol (HPP and NPP; see **f**. LC-MS traces are equally scaled overlays of total ion chromatograms (m/z 150-1500) recorded in negative ionization mode. These compounds were not detected by positive ionization analyses. **e,** Lipids co-purified by immuno-purification of flag-tagged CPSFL1 expressed in *E. coli* compared to purifications from control protein expressing cells. The differential display of 3 technical control LC-MS analyses (black, overlay) and 3 analyses of a lipid preparation copurified with flag-tagged CPSFL1 (red, underlay) identifies membrane lipids of *E. coli* and OPP (arrow). LC-MS traces are equally scaled overlays of total ion chromatograms (m/z 150-1500) recorded in negative ionization mode. **f,** Selected ion chromatograms from **e,** demonstrating presence of the molecular ions [M-H]^-^ of HPP, OPP, and NPP, namely m/z 569.4728 (C_41_H_61_O^-^; abundance x 1), m/z 637.5354 (C_46_H_69_O^-^; abundance x 1), and 705.5980 (C_51_H_77_O^-^; scaled abundance x 10). The 3 technical control LC-MS analyses do not contain these compounds. The 3 selected ion chromatograms (red) were extracted by expected monoisotopic masses with a mass tolerance of +/- 0.01 amu. The inserts show the measured m/z of [M-H]^-^ from HPP, OPP, and NPP, the mass differences between the compounds and the m/z deviation (Δ Da) compared to a prenyl-unit. Dashed vertical lines indicate the retention time of OPP.

However, the main dye accumulation resembled structures with comparable appearance as seen in TEM (Fig. 4a, TEM). Still, lipid droplets (LD) and vesicles are non-redundant entities but can be distinguished by the lipophilic core for LDs or enclosed hydrophilic cargo for vesicles (Fig.4a, scheme left). To verify an endocytosis-like mechanism, we tested internalisation of water-soluble and membrane-impermeable fluorescent dye, Tetrabromofluorescein (also called Eosin Y) from 1 mM to the growth medium according to^56^. Following expression of CPSFL1 and a control protein (C-terminus of KEA3) we measured endocytosis of dye by uptake from the culture medium and accumulation within the cells to confirm vesicular structures (Fig. 4b). Following 3 hours of protein expression, we first removed the dye from the outer layers of *E. coli* and surrounding media of the cells by repeated washing. The remaining dye was quantified spectroscopically. Only in *E. coli* cells expressing CPSFL1 the dye was retained (Fig. 4b, left). Furthermore, we localized the dye within the CPSFL1 induced structures rather than in the surrounding membranes by using fluorescence microscopy (Fig. 4b, middle). In addition, native purification of CPSFL1 also co-purified the endocytosed dye which was released upon solubilisation with the detergent Triton-X due to membrane rupture. (Fig. 4b, right). Thus, CPSFL1 mediates an endocytosis-like process when expressed in *E. coli* cells.

### Consequences of CPSFL1 expression on lipid and pigment metabolism in WT and engineered *E. coli* strains

Heterologous expression of CPSFL1 exhibits significant membrane transport activity. Thus, we analyzed the effects of CPSFL1 expression on the composition of the lipophilic fraction of *E. coli* cells. For this, we investigated the lipophilic phase of total cell extracts of CPSFL1-expressing and control strains (Fig. 4c). Following initial analysis by TLC we observed the strong accumulation of a single compound in CPSFL1 expressing cells. The substance was absent in WT cells (Fig. 4c, marked with *). In order to identify the substance, we performed LC-MS analysis (Fig. 4d and supplemental Figure 4). Interestingly, CPSFL1-expressing cells accumulated a lipophilic compound tentatively identified as octaprenylphenol (OPP)^57,58^(Fig. 4d). OPP identification was by match of exact masses of the molecular and adduct ions and by additional accompanying accumulation of analogous compounds with 7 or 9 prenyl units (Supplemental Figure 4). OPP is a membrane integral prenylquinone and an intermediate of ubiquinone biosynthesis^57^. Due to its hydrophobicity and metabolite channelling, OPP does not accumulate under normal conditions^19,58^. Instead, OPP accumulates only in mutants with defective downstream processing^59^; Thus, OPP accumulation in CPSFL1 expressing cells indicates a block in quinone biosynthesis downstream of OPP causing substrate accumulation of this intermediate. We suspected a direct binding of OPP to CPSFL1. Thus, we isolated recombinant CPSFL1 and analysed the co-purified lipophilic fraction by LC-MS (Fig. 4d, middle panel). Native CPSFL1 predominantly co-purified membrane lipids as reported above by using TLC (Fig. 3d and supplemental Fig. 3b). Low amounts of OPP could be detected in the CPSFL1 co-purified lipid fraction (Fig. 4d, lower right, supplemental Fig. 3b

### Tuning CPSFL1-mediated endocytosis in engineered *E. coli* via phosphatidylinositide biosynthesis and curvature modulation

*In vitro* experiments using GUVs supplemented with PPIs to the lipid mixture showed an increase in CPSFL1 membrane recognition, binding and deformation (Fig. 2). Here, we investigated the impact of curvature modulation by PPIs on CPSFL1-mediated endocytosis in *E. coli* (Fig.5). Since *E. coli* is naturally devoid of PPIs, we genetically engineered PPI synthesizing *E. coli* strains by introducing a plasmid expressing PI synthase (PIS), PI-4-kinase (PI4K) and PI4P-5-kinase (PI4P5K) for the sequential synthesis of PI, PI4P and PI(4,5)P_2_ within the plasma membrane of *E. coli* cells (Fig. 5a)^60^. The addition of *myo*-inositol into the growth medium induced PPI production^60^. TEM of cryofixed *E. coli* cells exhibited undulating inner and outer membranes with strong curvature probably due to PPI accumulation (Fig. 5b).

**Figure 5:**
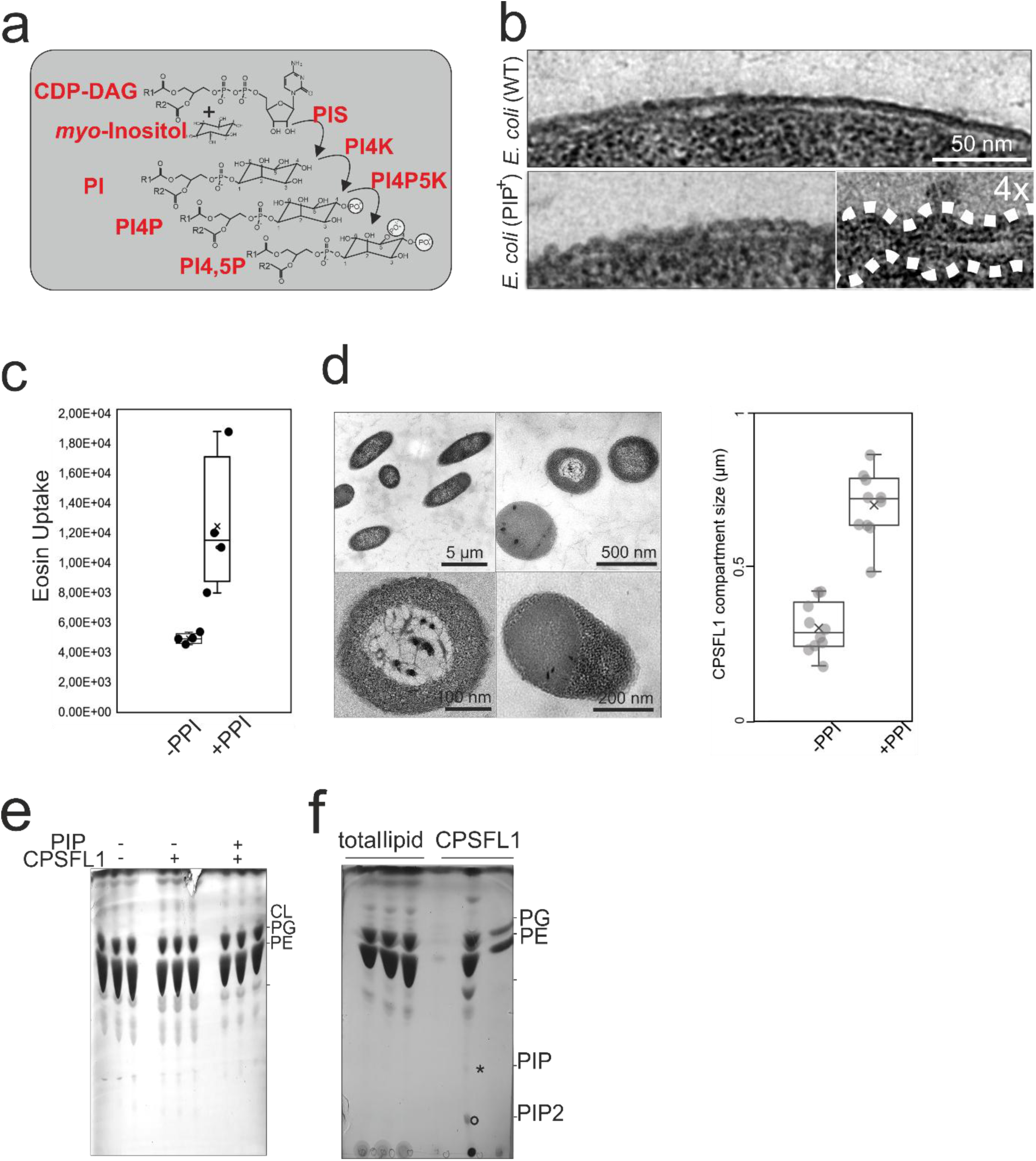
Regulation of CPSFL1 dependent vesicle formation via PPIs in genetically engineered bacteria. **a,** Engineered PPI Synthesis pathway introduced into *E. coli* cells. PIS: phosphatidylinositol-synthase, PIK: phosphatidylinositol-kinase, PI4P5K: phosphatidylinositol-4-phosphate-kinase. CDP-DAG: cytidine diphosphate diacylglycerol. **b,** Electron micrograph of *E. coli* cells expressing PPIs as compared to WT shows undulating outer and inner membranes probably due to curvature inducing properties of conical PPIs. **c,** Quantification of endocytosis as measured by dye uptake of cells expressing PPIs or PPIs and CPSFL1 in comparison to control strains. **d,** Electron micrograph of *E. coli* cells expressing PPIs and CPSFL1. Upon expression of CPSFL1 in PPI expressing cells, enormous electron opaque and electron dense structures appeared within the cells imaged using TEM. Analysis of the structures showed a double diamaterby equal cell size as compared to E. coli cells expressing CPSFL1 only. **e,** TLC of *E. coli* cells expressing PIPs or PIPs and CPSFL1 in comparison to total lipid extracts of WT *E. coli* cells. The synthesis of PI leads to the detection of an additional band in both PIP and PIP/CPSFL1 expressing lines. **f,** Comparison of lipid composition between total lipid extracts of PIP producing E. coli cells and purified CPSFL1. PPI levels stayed below the detection limit in total cell extracts but could be detected in purified CPSFL1 dependent vesicles.

The PPI producing strains were super transformed with a CPSFL1-FLAG and a control protein expressing plasmids respectively. To quantify the impact of PPI synthesis on CPSFL1 mediated endocytosis, we quantified again dye uptake from the surrounding medium as described above (Fig. 4b). In comparison to cells only expressing CPSFL1 a marked increase in endocytosis was observed (Fig. 5c). Next, the ultrastructure of the cells was analysed by TEM upon protein induction (Fig. 5d). Measurements of the lipophilic compartment of PPI expressing vs. CPSFL1-Flag only expressing cells showed a doubling in diameter of the osmiophilic compartment but constant cell diameter (Fig. 5d, right). Occasionally, white inclusions mimicking bigger vesicular structures without osmiophilic content were observed instead of dark compartments (Fig. 5d, lower left). Overall, single vesicles were rarely observed and the endogenous structures of large subcellular structures presumably formed by smaller vesicles. Thus, we performed a compositional analysis of the lipophilic fraction of WT, PPI and PPI/CPSFL1 co-expressing *E. coli* cells (Fig. 5e). Total lipid extracts showed no obvious changes of TLC stainable lipids (Fig. 5e). The likely low abundant PPIs could not be visualized by staining. However, the lipid profile of natively purified CPSFL1 from PPI expressing cells co-purified with a membrane lipid mixture enriched with substantial amounts of PI4P, and PI(4,5)P_2_ as indicated by lipid standards analysed by TLC (Fig. 5f, supplementary Figure 5a). PPI accumulation supports a directional transport of PPI containing membranes from the inner bacterial membrane by CPSFL1-mediated vesicle formation. In line with that, also carotenoid biosynthesis as a membrane associated process shows defects in *cpsfl1* mutant plastids^17^.

### Plant CPSFL1 co-purifies vesicular structures with chloroplast lipids

Previous results indicated a direct function of CPSFL1 in vesicular transport and identified a co-transport mechanism for lipids and prenylquinones in WT and PPIs in genetically engineered *E. coli* cells respectively. Intriguingly, plant mutants of CPSFL1 (*cpsfl1-1 or pitp7-1*) hint at a defect in the intervened prenylquinone metabolism in plastids by reduced β-carotin and plastoquinone levels^17,18^. Thus, we performed an orthogonal approach to *E. coli* cells with chloroplasts, to verify the lipid ligands of CPSFL1 *in vivo*. As observed in *E. coli* immunolocalization experiments on chloroplasts of transgenic *35S-CPSFL1-FLAG/cpsfl1-1* complementing and in *35S-CPSFL1-YFP* overexpression plants also detected similar distributions of gold particles within chloroplasts probed for CPSFL1-FLAG and CPSFL1-YFP proteins respectively (Fig. 6a). Next, we used CPSFL1-FLAG and fluorescent tagged CPSFL1-YFP lines for co-immunoprecipitation of native CPSFL1 from stromal preparations of osmotically ruptured chloroplasts (Fig. 6b). We verified the presence of CPSFL1-FLAG in immunoprecipitations by western blotting (Fig.6b, lower left). Next, we investigated CPSFL1-FLAG purifications by negative stain and electron microscopy (Fig.6 b, upper left). CPSFL1-YFP bound to magnetic nanobody conjugated beads could not be used for TEM imaging. In CPSFL1-FLAG globular structures were detected. In addition, these structures were absent in control experiments of co-immunoprecipitates from stromal preparations not containing tagged CPSFL1. Ultrastructural analysis of these structures identified a diameter of 33 nm (Fig.6d). To proof whether these structures could represent membrane vesicles, we analysed the lipophilic fraction by thin-layer chromatography and mass spectrometry (Fig. 6c). In comparison to control samples using a chloroplast localized GFP, we detected a CPSFL1 specific enrichment of chloroplast lipids (supplementary Fig. 6c, supplementary Fig. 6). This included a more than fivefold enrichment of the main membrane lipids of the inner envelope membrane and the thylakoids, namely, MGDG, DGDG, SQDG and PG. Whereas, an increased abundance was confirmed the composition of respective lipid subspecies and their c16/c18 fatty acid ratio remained unchanged (Fig. 6 c, lower panel). Furthermore, additional lipid species were detected (supplementary Fig. 6b). These included the typical lipid ligands of the SEC14 protein, PC and PI^61^. Another category includes signal lipids. These were tentatively identified by exact mass as potential acylated lipids, acyl-PG, acyl-DGDG and acyl-MGDG as well as the conical lipid DAG. The qualitative assessment of lipids indicates a CPSFL1 dependent enrichment of membranes. While mass spectrometric analysis did not allow for a relative molar composition of lipids within CPSFL1 preparations, low amounts of lipids did also not allow a clear visualization using TLC. However, the determined composition does not correspond to a typical chloroplast membrane^2,4^. This adds complexity to the idea of these structures as inner membrane vesicles (Fig. 6c, d). Alternatively, CPSFL1 might bind to two or more lipid types *in vivo* or in a sequential and dynamic process. Whether, PQ or a related molecule was copurified with membrane lipids remained unknown. Interestingly, ultrastructural analysis of *cpsfl1* mutants indicates changes in plastoglobule number and content and shows electron-transparent (non-osmiophilic, or white) thylakoid associated globular structures as compared to electron-opaque (osmiophilic, or black) plastoglobules observed in WT chloroplasts (Fig. 6e). In addition, *cpsfl1* mutant chloroplasts show an increased number of plastoglobules (Fig. 6f, left). However, plastoglobules of cpsfl1 were predominantly white as compared to black (osmiophilic) WT plastoglobules (Fig. 6f, right). This indicated a different composition of their core.

**Figure 6:**
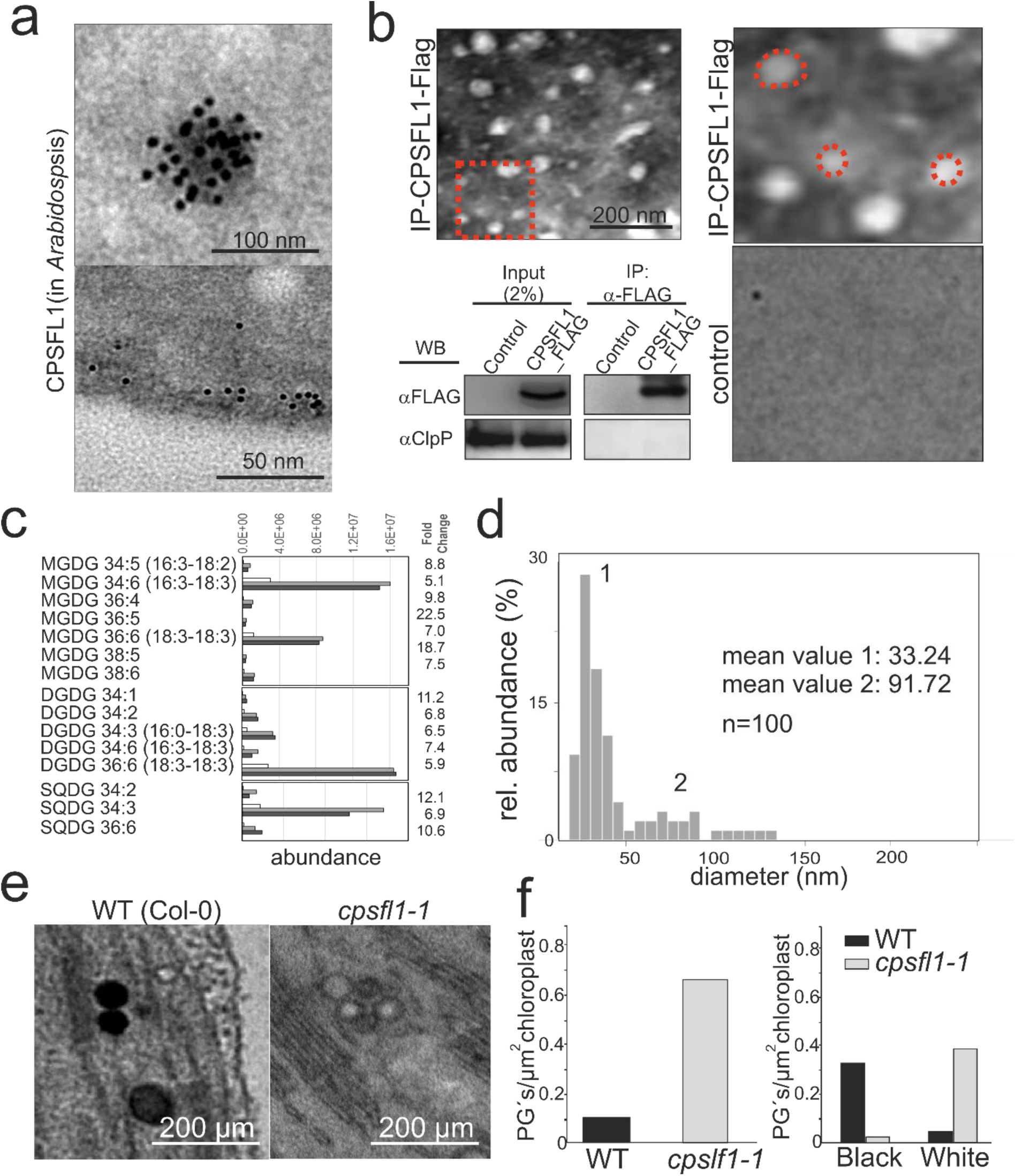
Native chloroplast CPSFL1 co-purifies membrane vesicles. **a,** Similar to *E. coli*, electron micrographs of immunogold labelled sections of chloroplasts from CPSFL1_Flag expressing plants show gold clusters at the envelope (lower panel) and within the stromal compartment (top panel). **b,** TEM micrographs of CPSFL1_Flag co-immunoprecipitates following negative staining show the presence of spherical structures (IP-CPSFL1-Flag). Red dotted square marks magnified region on the right. These structures were not observed in elutions from control protein C-term KEA3 (control, lower image). Immunoprecipitations were analyzed following SDS-PAGE using chloroplast as control by immunoblotting using FLAG specific antibodies, CPSFL1_Flag was shown to be immunopurified. ClpP was used as a negative control protein for Co-IPs and could not be detected. **c,** Characterization of the lipid fraction that was co-immuno-purified together with YFP-tagged CPSFL1 from pre-purified chloroplasts of respective genetically modified plants. Chloroplasts were pre-purified to avoid artificial contact of the fusion protein with eukaryotic membranes during the purification process. Non-targeted lipidomic analysis of the co-purified lipids revealed 261 mass features that accumulated at least 5-fold relative to a control lipid preparation from plants that express YFP targeted to chloroplasts. Selected mass features were among the top 1500 abundant (arbitrary units) of 3294 detected mass features. 130 of the selected mass features were annotated as molecular ions, adducts or *in source* fragments of lipid species from chloroplast-located lipid classes. Specific co-purified species of each lipid class are shown. Additional non-enriched lipid species of each class are omitted. Annotation was by match of predicted molecular mass and retention times to reference libraries or in the absence of reference compounds, namely Acyl-PGs and Acyl-MGDGs (Supplemental Figure S6), by match of predicted molecular masses. The plots show co-immunopurified abundances of each lipid species from the control preparation (white bar) and two independent co-purifications (grey and dark grey bars). Monogalactosyldiacylglycerols (MGDG), digalactosyldiacylglycerol (DGDG), sulfoquinovosyldiacylglycerols (SQDG). For further lipid classes, e.g., phosphatidylglycerol (PG), phosphatidylcholine (PC), chlorophylls, and acyl-MGDGs, refer to the supplement (Supplemental Figure S6). Fold changes to the right indicate the average enrichment of each lipid species across the two independent preparations compared to the control preparation. **d,** Size comparison of CPSFL1 bound vesicles isolated from *E. coli* and plants showed a significant difference. Whereas bacterial vesicles showed an average diameter of 15 nm, isolated plant vesicles with an average diameter of 40 nm were much bigger. **e,** Transmission electron micrographs of WT and *cpsfl1-1* mutant chloroplasts showing plastoglobules (arrows). Similar structures appeared close to thylakoids in *cpsfl1-1* mutants but appear pale. **f,** Quantification of plastoglobuli number (PGs) per µm^2^ in WT and *cpsfl1-1* mutant plastids. Quantification of electron-transparent (non-osmiophilic, or white) plastoglobules as compared to electron-opaque (osmiophilic, or black) plastoglobules observed in WT chloroplasts.

In summary, CPSFL1 emerges as a multifaceted protein influencing membrane dynamics, curvature modulation, vesicle formation, and endocytosis, giving a mechanistic framework for a vesicle mediated intermembrane metabolite co-transport for both bacteria and chloroplasts.

## Materials and Methods

### Plant cultivation and cloning

*Arabidopsis thaliana* WT (Col-0) and mutant plant seeds (CPSFL1-YFP, CPSFL1-Flag, cpsfl1-1) were germinated on 0.5× Murashige and Skoog^62^ agar medium enriched with 1% (w/v) sucrose. For experiments including *cspfl1-1* mutants, seedlings of all genotypes to be compared were grown on MS medium containing 1 % sucrose with 16-hour light (120 μmol m^−2^ s^−1^) at 22 °C, and 8 h dark at 22 °C for 4 weeks. *cpsfl1-1* mutant plants were identified using a PAM fluorimeter via decreased Fv/Fm values. For other experiments seedlings were transferred to soil and cultivated for 4 weeks under a diurnal cycle comprising 16 hours of light (120 μmol m^−2^ s^−1^) and 8 hours of darkness at 22 °C. CPSFL1-FLAG and *cpsfl1-1* plant lines were described previously^16^. CPSFL1-YFP was generated by cloning the cDNA of CPSFL1 omitting the stop codon into a linearized plasmid (pML74 was gift of Ralph Bock) carrying a C-terminal YFP under the control of the 35S promotor and the NOS terminator using infusion^®^ cloning. The constructs were introduced into *Arabidopsis* (Col-0) through a floral dip method by utilizing Agrobacterium tumefaciens (strain GV3101)^63^. Transgenic seeds were selected by Kanamycin resistance and CPSFL1-YFP expression was verified in rosette leaves microscopically. 35S GFP expressing plants were used as a control and obtained from Salma Balazadeh^64^.

### Bacteria strains, cultivation, plasmids and cloning

CPSFL1 was expressed in BL21 (DE3) pLysS cells. The plasmid for recombinant flag tagged CPSFL1 was described previously. Fluorescent tagged recombinant CPSFL1 variants were cloned into pET28a using infusion^®^ cloning. For that CPSFL1-YFP without the cTP was amplified using Primers…. and …. from the plant plasmid described above. For *recΔAH_(CPSFL1)_-YFP*, CPSFL1 was amplified using primers … and … from the same plasmid. For *recAH_(CPSFL1)_-YFP* the coding sequence of the amphiphilic helix was amplified using primers … and … from the cDNA sequence of CPSFL1 and cloned fused to YFP into pET28a by infusion cloning. A plasmid expressing the KEA3-C-terminus in pET28a was a kind gift from Ute Armbruster^66^. A Plasmid expressing Phosphatidylinosiltol-phosphate biosynthesis (p15aC-4D1D-5) was a gift from Sanford Simon (Addgene plasmid # 107866; http://n2t.net/addgene:107866; RRID:Addgene_107866)^60^. The plasmid encodes for Human phosphatidylinositol 4-phosphate 5-kinase type-1 α isoform 2 (PI4P5Kα, PI4P5K), Bos taurus phosphatidylinositol 4-kinase β (PI4Kβ, PI4K), Trypanosome brucei phosphatidyl inositol synthase (PIS), a chlorpamphenicol resistance and produces Phosphatidylinositolphosphates when introduced into E. coli (BL21 (DE3) pLysS) in medium supplemented with myo-inositol.

In order to co-express CPSFL1 or control proteins in PIP synthesizing cells we initially transformed E. coli cells (BL21(DE3) pLysS with p15aC-4D1D-5 plasmid and selected positive transformants via cAMP resistance. Cells were made chemically competent and were super-transformed with CPSFL1 plasmids. Co-expressing transformants were selected by double antibiotic selection against Kan and cAMP. For protein expression cells were grown in YT-medium until they reached an OD600 of 0.6-0.8. For expression of PIPs myo-inositol (5 µm final) was supplemented to the medium 1 hr before expression was induced. Protein expression was induced by addition of 0.5 mM IPTG and the culture was grown for additional 3 hrs at 30°C. Subsequent cells were harvested by centrifugation and pellets were immediately frozen in liquid nitrogen.

### Recombinant protein expression and purification

For the analysis of lipid ligands and ultrastructure of CPSFL1 particles, *E. coli* cells were thawn on ice, resuspended in lysis buffer and broken up by passing three times through a French press at 10.000 PSI @ 4°C. Subsequent protein purification was performed under native conditions by adding Ni-NTA (Quiagen) according to the manufactureŕs recommendation. In a last step buffer exchange of eluted protein solution was done against PBS using centricons (10.000 MW, Millipore) and proteins were diluted to 2 µg/µl, snap frozen in liquid nitrogen and stored at -80°C until use.

### Chloroplast isolation, plant protein extraction, Co-Immunoprecipitation, SDS-PAGE and Western blotting

Chloroplast isolation was done as described previously^11^. Chloroplast were ruptured by three cycles of freezing in N_2_ and thawing on ice. For Co-IP broken chloroplast were centrifuged at 14.000 x g for 20 min. at RT. The supernatant was incubated with Flag- or GFP-specific antibodies immobilized on agarose (EZview™ Red ANTI-FLAG® M2 Affinity Gel) or magnetic beads (Chromotek GFP-trap®) respectively. Following three washing steps in PBS and sedimentation by centrifugation at 1000xg for 1 min. at 4°C or magnetically. The resulting fraction was used for negative staining or lipid extraction or SDS-PAGE.

### Lipidomic Analyses

Relative changes of lipid abundances were analysed by lipidomic profiling using liquid chromatography-mass spectrometry (LC-MS) as was described by ^65^ with modifications reported by ^66^. Samples of *E. coli* cells BL21 (DE3) pLysS in logarithmic growth phase were harvested by centrifugation and snap-frozen in liquid N_2_. Cells expressing flag tagged CPSFL1 were compared to *E. coli* cells expressing the C-terminus of KEA3. Approximately approximately 200 mg fresh weight of cell pellets or of flag-tagged CPSFL1 protein that was his tag-purified as described above from an approximately similar amount of *E. coli* cells or YFP-tagged CPSFL1 protein or GFP (control) that was immuno-purified from pre-purified chloroplasts (see Co-immunoprecipitation in methods description) were extracted by a Bligh and Dyer based method according to^67,68^ with modifications. In brief, samples were mixed with chloroform/methanol/HCl (50:100:1, v/v). Following centrifugation (10 000.g) for 2 min, supernatant was used to induce a two-phase system by the addition of 1 volume of chloroform and 0.8 volume of 0.9% (w/v) NaCl. Samples were mix rigorously and centrifuged for 2 min to get the two-phase system. The upper-phase was discarded and the organic lower phase was washed three times with H_2_0/methanol/HCl (50:50:1, v/v). Lipid extracts were dried under a stream of nitrogen and stored at -80°C until further use.

Re-dissolving of lipids into 150 μL acetonitrile:isopropanol (7:3, v/v, UPLC grade; BioSolve) with 1% (v) 1 M NH_4_Ac and 0.1% (v) acetic acid and subsequent ultra-performance liquid chromatography (UPLC) of 2 μL re-dissolved lipid samples by an Acquity UPLC system with an BEH C8 reversed-phase column (100 mm × 2.1 mm with 1.7 μm particles; Waters GmbH, Eschborn, Germany, http://www.waters.com), as well as mass spectral analyses by an orbitrap mass spectrometer (Exactive, Thermo Fisher, Waltham, USA, http://www.thermofisher.com) was exactly as described previously ^69^). Each sample was measured by both, positive- and negative-ionization mode. Data processing of chromatogram files included baseline correction, chemical noise subtraction, alignment, and peak detection was by REFINER MS 5.3 software, https://www.genedata.com/, according to^69^. Mass features characterized by the 2 orthogonal parameters, mass-to-charge ratio (m/z) and retention time (RT), were assembled into a numerical data matrix with respective abundances (arbitrary units) of each sample. Lipids were annotated by matching to reference libraries of expected m/z and RT values^65,66^. This method distinguishes lipid classes and lipid species according to the sum of carbon atoms and the degree of unsaturation of their fatty acids. We name lipids accordingly, e.g. PC 34:0 for 1,2-diheptadecanoyl-sn-glycero-3-phosphocholine. Lipid isomers are chromatographically resolved and named by extensions, e.g., (1) or (2) in chromatographic order. To characterize the acyl-chain composition of monogalactosyldiacylglycerols (MGDG) and digalactosyldiacylglycerols (DGDG), we co-analysed commercially available authenticated preparations, namely, synthetic MGDG 18:1-18:1 (Sigma-Aldrich/Avanti Polar Lipids, 840531P), MGDG 18:2-18:2 (Sigma-Aldrich/Avanti Polar Lipids, 840532P), and MGDG 18:3-18:3 (Sigma-Aldrich/Avanti Polar Lipids, 840533P), and defined biological mixtures of MGDGs (Sigma-Aldrich/Avanti Polar Lipids, 840523P) and DGDGs (Sigma-Aldrich/Avanti Polar Lipids, 840524P), according to the manufacturer’s analysis certificates. In these cases, we use name extensions with acyl chain designations, e.g. PC 34:0 (PC 17:0-17:0). The *sn*-positions of the fatty acids within the glycerol-lipid species are not resolved. Annotations of prenylphenols were supported by exact monoisotopic mass to molecular formula matching and molecular formula to structure searches at https://www.chemcalc.org/mf-finder, https://www.metabolomicsworkbench.org, https://www.genome.jp/kegg/compound/, https://biocyc.org/cpd-search.shtml, and https://pubchem.ncbi.nlm.nih.gov.

For lipid abundance analyses we used approximately equal sample amounts and performed background subtraction using non-sample controls. We report abundances in arbitrary detector units. For qualitative analyses we selected lipids that were at least 8-fold (*E. coli*) or 5-fold (*Arabidopsis thaliana*) more abundant relative to respective control samples and were among the top 1500 (*E. coli*) and top 1000 (*Arabidopsis thaliana*), most abundant within each experiment.

### Lipid extraction and analysis by TLC

TLC was performed according to Munnik et al ^70^. In brief, Silica60 glass plates were impregnated by dipping into a solution containing 5 mM oxalic acid 2 mM EDTA in 40% methanol. Subsequent plates were activated by baking at 120°C for at least 30 min. Plates were pre-run using acetone, dried and used for TLC. Samples were separated using alkaline solvent [(Chloroform: Methanol: Ammonimhydroxid: ddH20) 90:70:4:16)], dried and immersed in a solution containing 10% Copper(II)sulphate in 8% phosphoric acid. Lipids were visualized by charring at 180 °C for 10 min^71^. Lipids were quantified by intensity of the bands using ImageJ.

### Spectroscopic analysis

For spectroscopic analysis solutions were either used directly or pigments were extracted by dilution with pure acetone (1:5 dilution). Following centrifugation supernatants were measured using a spectrophotometer. For Eosin (Sigma) absorbance was measured at 700 nm.

### Confocal microscopy

GUVs were either imaged on an inverted spinning disc (custom build microscope) or on an upright confocal microscope (Leica Systems, Wetzlar, SP8) For this GUVs were deposited on a BSA (1 mg/ml) coated coverslip first and allowed to sediment for 10 min. at RT before imaging. All multichannel imaging was recorded in sequential mode. YFP was excited at 514nm using an Argon laser and emission was recorded between 520 and 560 nm. DiI was excited at 561 nm and fluorescence emission was recorded between 577 and 670 nm. Eosin was excited at 561 nm and emission was recorded between 653 and 695 BODIPY (Thermo) was excited at 488 nm and fluorescence emission was recorded between 505 and 586nm, FM4-64 (Thermo) was excited at 633 nm and fluorescence emission was recorded between 676 and 769 nm.

### Transmission electron microscopy and sample preparation

#### Negative staining

For negative staining a drop (5-10 µl) of sample was applied to the surface of a carbon coated formvar grid (Nickel, 200 mesh) and allowed to partially dry. Afterwards grids were washed three times with water to remove salts and phosphates and a drop of aqueous Uranylacetate (1%) was applied to the grid and incubated for 30 sec. following three rinses with ddH2O grids were rinsed three times with ddH2O dried and used for imaging.

#### High-pressure freezing and freeze substitution

*E. coli* samples were high-pressure–frozen using a high-pressure freezer (Leica HPM100). Subsequent samples were either freeze-substituted in 1% OsO4 and 0.1% uranyl acetate in acetone for ultrastructural analysis or in 0.5% uranyl acetate in acetone for immunolabelling and embedded into LR White medium at −20°C and polymerized at −20°C using UV light.

#### Immunogold-labeling

Immunogold labelling was done on cryo-fixed and freeze-substituted 100-nm-thin sections of samples embedded into LR White. Initially samples were incubated for 1 hour at room temperature in blocking buffer [phosphate-buffered saline–Tween 20 (PBST) containing 2% bovine serum albumin and 0.1% fish gelatin (Sigma-Aldrich)] to reduce unspecific binding. Subsequent antigens were immunodecorated by incubation in blocking buffer containing anti-Flag (1:100 dilution, mouse; Sigma) antibodies for 1 hour at room temperature. Following six rinses with PBST buffer for 3 min each, primary antibodies were detected by incubation Sigma-Aldrich] in blocking buffer containing 20 nm colloidal gold labelled secondary antibodies (for mouse; Cell Signaling Technology, Danvers, MA) for 1 hr at RT. Afterwards unbound antibodies were removed with six rinses in PBST and three rinses with double-distilled water for 3 min each and samples were contrasted as described above.

#### Imaging

For TEM analysis, sections were cut using a Leica UC-6 ultramicrotome. Contrasting of sections was done using methanolic uranyl acetate (2% in 50% methanol) for 30 min, followed by a 10-min incubation in lead citrate (Reynolds’ stain). Images were either acquired with a Zeiss EM 912 Omega TEM (Carl Zeiss, Oberkochen, Germany) or a JEOL JEM F 200 (JEOL, Germany).

### SUV and GUV production and sedimentation assay

Liposomes (SUVs) were prepared using thin-film rehydration of phospholipid mixtures and extrusion. Synthetic lipids (10 mg from chloroform stock solutions) at a mass ratio of 1: 1 DOPC: DOPG were mixed in a 15 mL glass and chloroform was removed using a gentle stream of N_2_(g). To remove residual solvent, vials were placed in a vacuum desiccator for 1 h. The dried film was allowed to rehydrate for at least 30 min at RT by adding 1 mL of sucrose containing PBS with a total osmolarity of 200 mOsm (measured with freezing point osmometer Osmomat 3000, Gonotec) and vortexed periodically. The resulting solution was then extruded using a hand-held Mini-Extruder (Avanti) equipped with a 30, 50, 100, 200 or 400 nm polycarbonate membrane (Whatman). 20 passes through the membrane were performed to guarantee monodisperse size distributions which was determined by DLS measurements using a Zetasizer NanoZS (Malvern Instruments Ltd.). For binding assays, proteins in PBS containing glucose at the same osmolarity were added to the SUVs. Following incubation at RT for 30 min, samples were centrifuged at 100xg for 5 min. Supernatant and pellets were used for biochemical analysis and pellets were used for imaging.

GUVs were prepared by polyvinyl alcohol (PVA) assisted swelling in PBS buffer^72^. For this a 5% (w/w) solution of PVA (with MW 145000, Sigma) was prepared by stirring PVA in water while heating at 90°C. PVA-coated substrates were prepared by spreading 100–300 *μ*L of PVA solution on a microscope slide. The slide was then dried for 30 min in an oven at 50°C. Afterwards 10–20 *μ*L of lipid mixtures (2-4 mM) dissolved in chloroform were spread on the dried PVA film and placed under vacuum for 30 min to evaporate the solvent. A chamber was formed using a custom-made Teflon frame and a cover glass and filled with sucrose or sucrose containing PBS buffer (200 mOsm). Following 15 min. incubation, GUVs were transferred into an Eppendorf tube using a pipette until further use. For protein binding studies proteins were added in PBS containing glucose in equal osmolarity in a final concentration of 0.5 µM.

### Dye uptake and endocytosis assay

Experimental setup was performed according to ^56^. For this, *E. coli* cells expressing either CPSFL1-Flag, CPSFL1-Flag and PPIs, control plasmid or control plasmids and PPIs were grown in YT medium at 37°C in the presence of Kan or Kan and cAMP respectively. One hour prior to induction of protein expression, fluorescent test molecules 5-(6)carboxyfluorescein (CF, Molekula, Gillingham, UK,10 mM final) and Tetrabromofluorescein (Sigma-Aldrich, Sydney, Australia, 20 mM final were added as solids directly to the growth medium. For PPI co-expression myo-inositol was added at 5 µm final concentration. FM4-64 or BODIPY were added to the cell suspensions in 2-10 µM final concentration. Protein expression was induced by the addition of isopropyl thiogalactoside (IPTG (Roth), 500 µM final) when an OD of 0,6-0,8 was reached. Protein expression was allowed to progress at 30°C for three more hours. Cells were harvested for analysis by centrifugation and washed for the removal of unincorporated fluorescent dyes and dyes trapped in the periplasmic space in cold Tris-buffered saline pH 7.5. Re-suspension and sedimentation were repeated until no fluorescence was detectable in the culture supernatant. For quantitation of dye uptake, the optical density of re-suspended cells was measured at 700 nm.

## Discussion

In summary, we identified a CPSFL1 dependent mechanism of membrane vesicle transport by its ability to sense, stabilize and promote membrane curvature (Fig.1b-d, Fig. 2d, Fig. 5c, Fig. 6). To date, multiple naturally occurring mechanisms eventually leading to membrane curvature have been described^27,41,54,73,74^. This also includes lipid composition of membranes^36,37^. Lipids like PA and PPI can act as curvature inducing lipids and are recognized by CPSFL1 with high specificity^16,17,75^. Recently identified chloroplast localized SEC14 homologs 5 and 7 have been shown to participate in the intermembrane transport of PA mediated by the TGD complex^76^. In line with that, synthetic membranes with typical chloroplast lipid compositions, containing PPI were also targeted and reshaped by CPSFL1 with high specificity (Fig. 1b-e, Fig. 2a). Interestingly, PPIs and PA are both, negatively charged and conical. In contrast, acyl-MGDG is uncharged and characterized by fatty acids attached to the head group of MGDG^77^. This may also affect lipid geometry, hydrophobicity, and charge and could potentially influence bilayer stability and induce membrane curvature. Another crucial feature in natural membranes is lipid sideness^78,79^. This is particularly caused by compositional (head or acyl group) and/or physical (lipid packing order, charge, hydration and H-bonding) between the inner and outer leaflets of lipid bilayers^79^. Packing defects or trans bilayer lipid asymmetry, lead to induction of spontaneous curvature^74^. According to preparation methods synthetic systems also GUVs can exhibit trans bilayer asymmetries^80^. Interestingly, addition of membrane binding proteins or altering transmembrane ion composition can induce these packing defects and curvature^81^. Binding of CPSFL1 to small vesicles with increased membrane curvature or packing defects supports these interpretations and highlights a direct function of CPSFL1 in vesicle traffic (Fig. 1d-e). Co-purification of CPSFL1 together with vesicular structures from bacterial and chloroplast extracts further supports direct interaction of vesicles with CPSFL1 (Fig. 3c-e, 4b, 5d, 6b). In line with that, CPSFL1 also binds to GUVs with chloroplast lipid composition largely independent of the presence of specific lipid species like PPI or PA (Fig. 2). In addition, these GUV experiments and ultrastructural analysis showed that CPSFL1 functions in membrane deformation (Fig. 2 1d,2e, 3c, 6b). Surprisingly, expression of CPSFL1 in a prokaryote leads to the formation of cytoplasmic vesicles as shown by electron microscopy linking CPSFL1 function to vesicle formation. (Fig. 3a, c). While E. coli does not naturally engage in endocytosis or form internal vesicles under standard conditions, certain experimental manipulations can induce the formation of vesicle-like structures within these bacteria^82^. Known mechanisms required for membrane deformation, invagination and vesicle fission are insertion of an amphiphilic loop and helix, intrinsically curved protein scaffolds (like BAR and ESCRTIII), lipid flipases or multidomain scaffolds^27,32,41,45,53,73,74,83–86^. Analysis of the CPSFL1 purified vesicle protein composition identified almost exclusively CPSFL1 protein (Fig. 3d). We characterized an amphiphilic helix located at the n-terminal region of the CRAL/TRIO domain of CPSFL1 required for membrane tethering (Fig.1a and b). While CPSFL1 is anchored to membranes by its amphiphilic helix, its soluble part formed by the majority of the CRAL/TRIO domain protrudes from the membrane (Fig.1a). Intrinsically disordered Sec14 proteins (IDPs) induce membrane deformation due to compressive stress during liquid-liquid phase separation^86–89^. Steric repulsion between proteins on biological membranes is known as a mechanism responsible for membrane re-shaping and eventually fission^90^. To amplify steric pressure, (i) hydrophobic insertions must anchor proteins strongly to the membrane surface and (ii) proteins need to be bound to the membrane in a high coverage^86^. Immunogold studies on *E. coli* and plastid CPSFL1 indicates oligomerisation of CPSFL1 proteins on vesicles or budding membranes *in vivo* (Fig. 3b, 6a). Whether vesicles result via a multidomain assembly like in clathrin-coated vesicles remains unclear^33^. Protein crowding and repulsion on their surface may prevent fusion of CPSFL1 bound vesicles into bigger structures (Fig. 3b, 6a). In a more detailed investigation using tracking of membrane-bound endocytosis dyes like FM4-64 and soluble membrane impermeable fluorescent dyes in the model prokaryote *E. coli* we highlighted an endocytosis-like transport activity upon expression of CPSFL1 (Fig. 4a, b). In fact, many Sec14 proteins have been assigned to function in vesicle traffic. However, no direct function had been shown for CPSFL1 so far. Thus, transport of PI to Golgi membranes by founder Sec14 is used to produce the signalling lipid phosphatidylinositol-phosphate (PI4P). PPIs in turn recruit proteins for vesicular transport between the *trans* Golgi network and the plasma membrane^24^. Thus, *sec14* mutants show a defect in vesicle transport due to low Golgi PI and PIP levels^91,92^.

This raises the question about the function of the CRAL/TRIO domain of CPSFL1. For the yeast Sec14 protein, lipid transfer activity and membrane binding are both promoted by membrane curvature^26^. We showed that CPSFL1s CRAL/TRIO domain is needed to recognize membranes with strong curvature (Fig.1b). Expression of CPSFL1 in *E. coli* identified its impact on quinone biosynthesis and highlighted an interference with the native quinone biosynthesis pathway “down-stream” of the prenylquinones possible quinone transport mechanism (Fig. 4d, e). However, our experiments rather exclude a direct binding of prenylquinones by CPSFL1 (Fig. 3d, native extracts). Instead, we interpret prenylquinones and carotenoids rather as cargo within the vesicle membranes (Fig. 3d, 4d, supplemental Fig. 5). This corresponds well with the phenotype of *cpsfl1* mutants, that show reduced levels of both, carotenoids and quinones^17,18^. Their biosynthesis pathways both feed on geranylgeranyl pyrophosphate (GGPP) a common precursor synthesized in the chloroplast envelope membrane^7,93^. Both pathways require also transport of the final products to the thylakoid membranes^7,19,93^. Surprisingly, the intermediate, OPP, and not the final product Ubiquinone accumulated in *E. coli* cells (Fig.4d). A conclusive explanation came from the analysis of the bacterial Ubi synthesis pathway^94^. OPP is the last membrane integral intermediate in Ubiquinone biosynthesis. Subsequent quinone biosynthesis occurs via a soluble metabolome encoded by UbiE-K^58^. Consequently, OPP needs to be exported from the membranes. How OPP leaves the membrane is completely unknown^58^. Flag-tagged CPSFL1 co-purifies membrane lipids of *E. coli* in agreement with the assumption that CPSFL1 binds to membranes of *E. coli* and may be involved in vesicle formation and endocytosis. Whether the observed copurification is caused by direct interaction of CPSFL1 with OPP or whether OPP is copurified as part of the membrane fraction that binds to OPP is unclear. These CPSFL1 bound membranes contain prenylphenols as cargo, but we cannot rule out that this observation is non-specific and caused by general cellular over-accumulation of prenylphenols. *E. coli* appears to store the accumulated OPP, HPP and NPP in a compartment, e.g. a lipid droplet that may have CPSFL1 bound at the surface or in a membrane to which CPSFL1 does not preferentially bind.

Another possible explanation is that accumulation of lipophilic substances like polyisoprenoids (e.g. carotenoids or quinones) or triglycerids between bilayer leaflets creates packing stress ^43,95^. Also phase transitions of the non-bilayer lipids like MGDG in plastids and PE in bacteria respectively or Acyl-MGDG could induce packing stress. This could be sensed and resolved by CPSFL1 via membrane vesiculation even in the absence of signalling lipids. PPI expressing *E. coli* cells also showed increased endocytosis presumably due to increasing the amount of cone shaped lipid, like PIP^41^.

While the mechanisms of eukaryotic vesicular traffic are widely understood, chloroplast vesicle transport represents a long-standing conundrum^21,96–98^. Whether stromal vesicles are actually trafficking between the envelope and thylakoid membranes is also still unclear^19^. Still, chloroplast vesicles are absent upon loss of CPSFL1 and return upon CPSFL1 overexpression^99,100,16^. A similar phenotype is described for the plastid protein VIPP1^23^. Considering thylakoid membrane biogenesis, CPSFL1s function as phosphatidylinositol transfer protein (PITP) might be highly relevant for target protein recruitment^34,101,102^. The chloroplast protein Vipp1^34,84^ concentrates in curved membranes and shapes thylakoids^84,102,103^. In addition, VIPP1 is a member of the ESCRTIII family of membrane deforming proteins^35,102,104,105^ shows a high affinity for PIPs ^34^. *E. coli* also encodes for a homologue of VIPP1, termed Phage Shock Protein A (PspA)^104,106^. However, in comparison to VIPP1, PspA lacks the c-terminal domain which contains the PIP binding pocket required for functional complementation VIPP1 mutants^107^. Whether eukaryotic PPIs actually exist within plastid membranes is still unclear. Our lipid analysis did not allow for their detection (Fig.6). For this, other methods like *in vivo* radiolabelling prior to immunoprecipitation or immunologic detection in fat blot assays have to be applied in future experiments^68,108^.

Many Sec14 proteins transiently interact with membranes as lipid transfer protein (LTP) to deposit, extract or exchange lipids using their lipid binding domain (LBD), encoded in the cellular retinaldehyde-binding protein (CRALBP) and TRIO guanine exchange factor domain (CRAL/TRIO)^109,110^. The founder protein, Sec14, and many others transport PI in exchange to PC^24^. These proteins, classified as phosphatidylinositol-transfer proteins (PITPs) locally concentrate a lipophilic substrate or promote and stabilize membrane contact sites using their LBD^110,111^. Altered membrane lipid composition can directly influence membrane structure and properties or serves as signal for effector protein recruitment^53–62,80^. The key to understand the molecular function of Sec14 proteins thus lies in the identification of the in vivo lipid ligands. Here we present a functional characterization of chloroplast localized SEC14-like protein 1 (CPSFL1) including lipidomic analysis of bacterial and native *in vivo* lipid ligands of CPSFL1, which were isolated from chloroplasts by immunoprecipitation, revealed, among others, PC and also PI (Fig. supplementary Fig. 6b). In contrast to all other membrane systems phospholipids are highly underrepresented in the internal membranes of the chloroplast^2,4^. PC as a major phospholipid in most of the membranes within a plant cell, can barely be detected in internal membranes of the chloroplast. PI which is also a major phospholipid in eukaryotic membranes is a minor constituent of inner envelope and thylakoid membranes^4^. Thus, analogous lipid binding of CPSFL1 to cytosolic SEC14 within the chloroplast is conceivable. Specific enrichment of underrepresented lipids from soluble chloroplast extracts supports a specific lipid transport activity of CPSFL1 in plastids (Fig. 6). Furthermore, expressed in the cytoplasm of yeast cells CPSFL1 complements *sec14* mutants^16^. Similar results were also obtained from PPI producing *E. coli* strains which expressed CPSFL1 (Fig. 5). Following induction of PPI synthesis using myo-inositol, PPIs were barely detectable in total lipid extracts (Fig. 5e). In contrast, native CPSFL1 purifications contained stainable amounts PPIs indicating a PITP function of CPSFL1 (Fig. 5f). PPI transport is also conceivable to complement yeast SEC14 mutation^16,112^.

An unusual candidate enriched in lipid fractions of CPSFL1 natively co-purified from the chloroplast stroma is acyl-MGDG (supplementary Fig. 6b). Acyl-MGDGs are classified as oxylipins or Arabidopsides^77^. They serve as signal molecules in the regulation of developmental processes, plant stress response, and innate immunity^77,113^. Interestingly, several proteins with Sec14 domains were copurified with subfractions containing MGDG acyl transferase activity^77^. Acyl-MGDG and related Arabidopsides were frequently detected in experiments using wounding and chilling stress^77,113,114^. Under these conditions increased numbers of chloroplast vesicles have been shown^21,115,116^. Whether acyl-MGDG and vesicle formation are linked is unclear. Mutant plants lacking acyl-MGDG show no effect in chloroplast biogenesis^117^.

Since these lipid species, are rather low abundant in chloroplast lipid profiles but purified together with a mixture of the main membrane lipids using CPSFL1, it remains unclear whether PC, PI or acyl-MGDG are directly bound by CPSFL1. The main lipid composition corresponded to the inner membranes of chloroplasts and thylakoids in plants or the inner bacterial membranes in *E. coli* (Fig. 3e, 6c). Consequently, our results correspond to a mixture of CPSFL1 proteins with single lipid transport and CPSFL1-bound membrane fragments with packing defects or high curvature. Alternatively, we identified for the first time the lipid stoichiometry of bacterial or chloroplast membrane transport vesicles bound by CPSFL1 (Fig. 3e and 6c, supplementary Fig. 6b). Both the lipid compositions of bacterial and plastid membranes as of their respective vesicles purified using CPSFL1 are very different. Of course, the diversity of lipid species differs between the two organisms. This could also explain the differences in the size of the respective purified vesicles (Fig. 3c, 6d).

Dramatically decreased PQ content of *cpsfl1-1* mutants could point towards a role in quinone transport ^18^. However, *cpsfl1* mutants also have reduced numbers of plastids/cell and less thylakoids/chloroplast. To understand, whether this could be caused by a proportional lower thylakoid membrane abundance or a defect in transport we investigated the ultrastructural phenotype of *cpsfl1* mutants. In fact, less thylakoids in reduced numbers of chloroplasts/cell could account this reduction.

## Acknowledgments

We thank the Deutsche Forschungsgemeinschaft (DFG) and the Max Planck Society (MPG) for funding. We thank Prof. Dr. Ralph Bock and his team from the Max Planck Institute of Molecular Plant Physiology (MPIMP, Potsdam, Germany) for supporting and hosting the project and engaging in insightful discussions.

We thank Dr. Nadja Tarakina and Bolortuya Badamdorj from the Max Planck Institute of colloids and interfaces for providing access and excellent technical assistance to the transmission electron microscope.

We thank Prof. Petra Bauer at the Institute of Botany, at the Heinrich Heine University (Düsseldorf, Germany) for hosting the project and discussions.

## Author contributions

AH conceived the study. AH, MS and AE performed the experiments. AH, MS, SP, RD, AE and JK designed experiments and analyzed the data. AH wrote the manuscript with the help of all authors. All authors agreed to the submission of the manuscript.

