## Supplemental Information for "Insights into CPSFL1 Induced Membrane Dynamics: A Multifaceted Regulator Linking Vesicle Formation to Thylakoid Biogenesis"

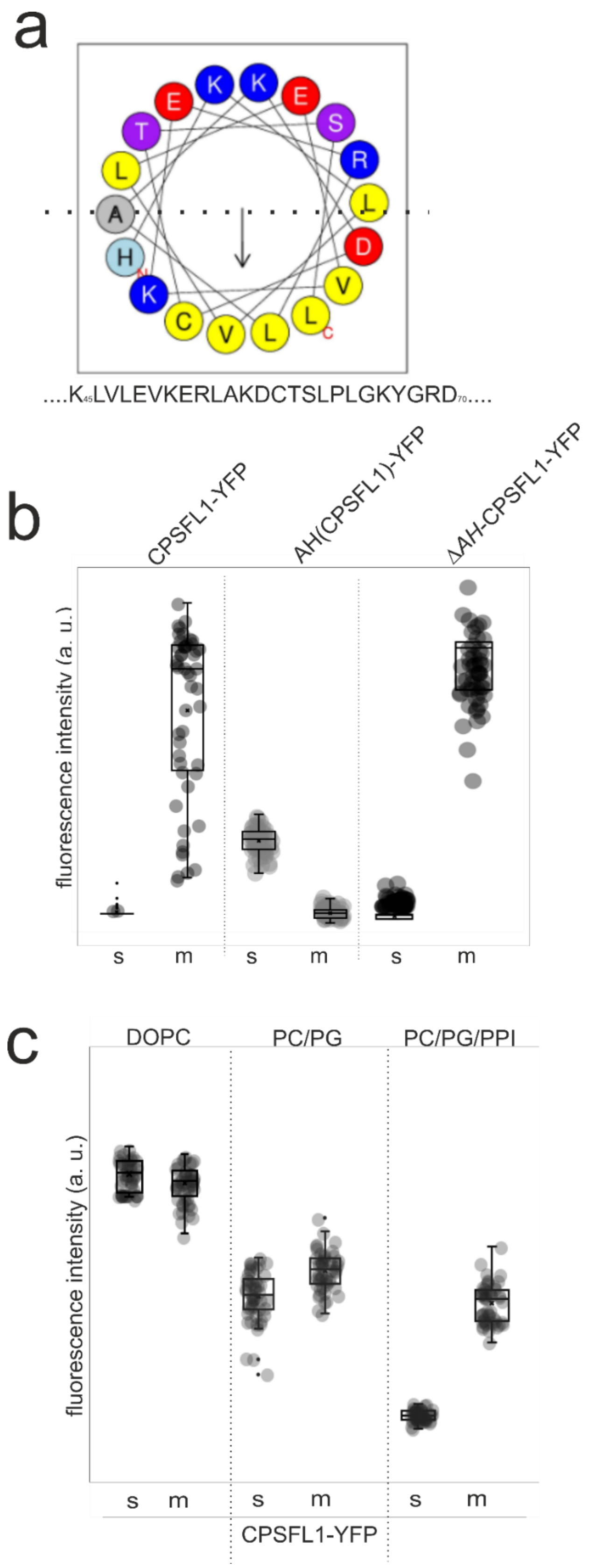


**Supplementary Figure 1**

**a,** *In silico* prediction of the CPSFL1 using HeliQuest predicted the CPSFL1 $\alpha$-helix between aa$^{45}$ and aa$^{70}$ through partitioning of hydrophobic and polar residues as amphiphilic. The arrow in helical wheels corresponds to the hydrophobic moment. Configurations and helical wheel representations are colour coded for residues: yellow for hydrophobic, purple for Ser (S) and Thr (T), blue for Lys (K) and Arg (R), red for acidic, grey for small residues (Ala, A and Gly, G), and light blue for His (H).

**b,** Quantification of confocal microscopy data on recombinant YFP tagged CPSFL1 and mutant variants with GUVs. Following addition of YFP tagged recombinant CPSFL1 protein variants (CPSFL1-YFP, *CPSFL1-ΔAH-YFP*, $AH_{(CPSFL1)}$-YFP) to GUV suspensions (1 μM final), membrane bound (m) and unbound GUV surrounding (s) protein fluorescence (YFP) was quantified. n=50.

**c,** Quantification of confocal microscopy experiments using recombinant fluorescent CPSFL1-YFP (1μM final) and GUVs neutral and charged and conical curvature inducing phospholipid containing GUVs. Protein fluorescence (YFP) and GUV fluorescence (DiI) was imaged and overlayed (merged) using confocal microscopy. Membrane bound (m) and unbound GUV surrounding (s) protein fluorescence (YFP) was quantified. n=50.

a

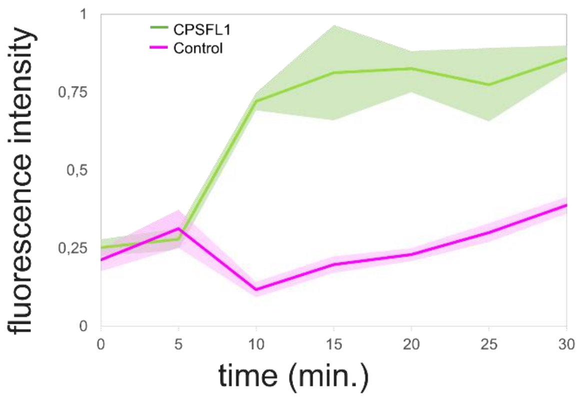


b

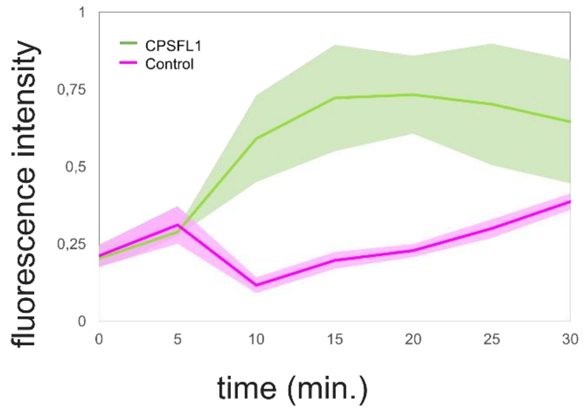


**Supplementary Figure 2**

**a,** Quantification of lipid fluorescence over time. Following CPSFL1-YFP addition, GUV deformation was accompanied by decrease in the fluorescence of the membranes and an increase in the fluorescence of the background (green). In control experiments fluorescence intensities remained unchanged (magenta).

**b,** Quantification of lipid fluorescence over time. Addition of CPSFL1-YFP to GUVs containing PI4P was accompanied by decrease in the fluorescence of the membranes and an increase in the fluorescence of the background (green). In control experiments fluorescence intensities remained unchanged (magenta). Fluorescence was quantified using ImageJ by taking multiple ROIs (n=15) of equal size on membrane and surrounding regions for each time point.

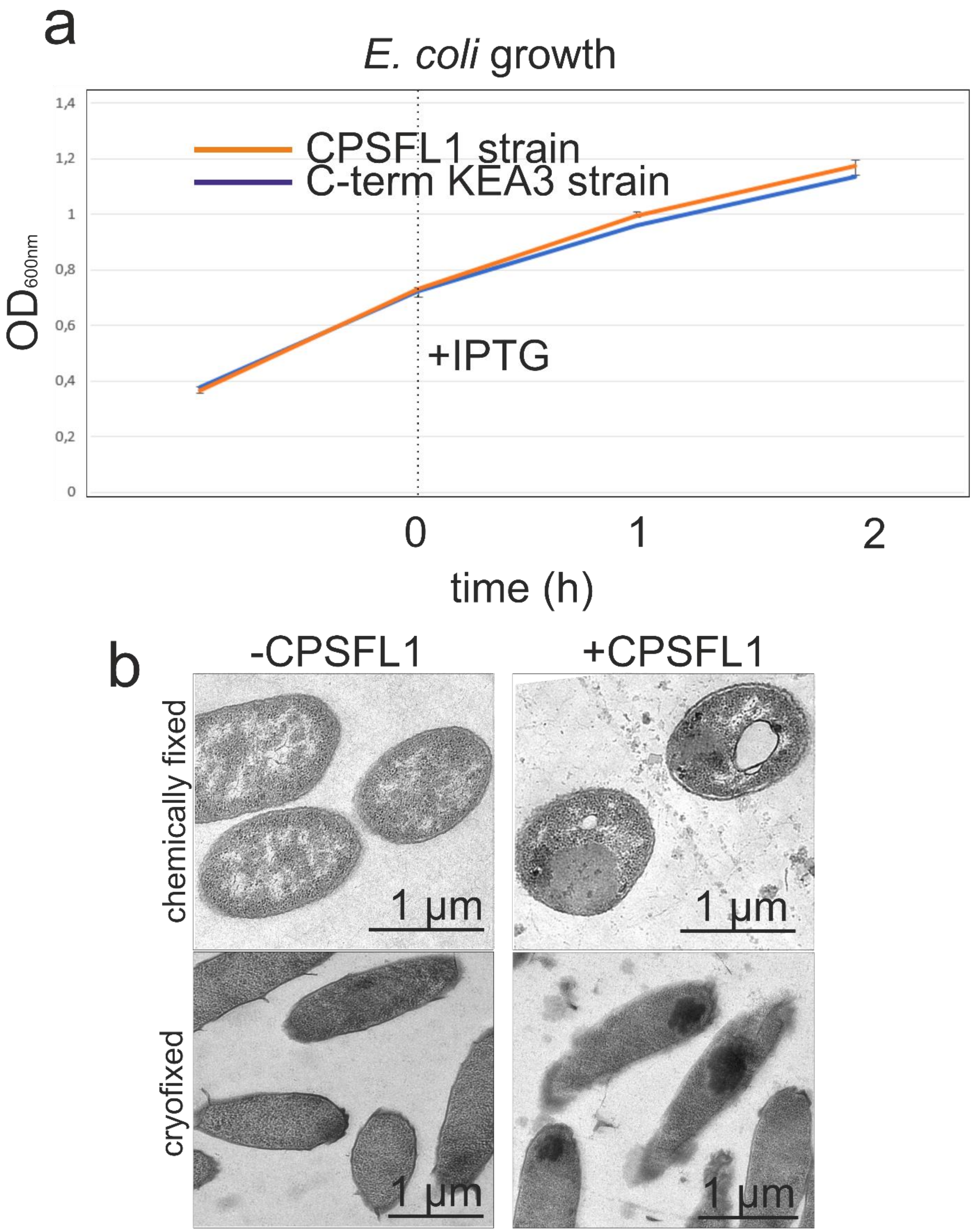


**Supplementary Figure 3**

**a,** Growth curves of *E. coli* cells expressing CPSFL1-Flag or the control protein (C-terminus KEA3) before and after addition of IPTG. No differences were observed.

**b,** Transmission electron microscopy (TEM) images of *E. coli* cells expressing a control protein (-CPSFL1) or CPSFL1 (+CPSFL1) following chemical or cryofixation.

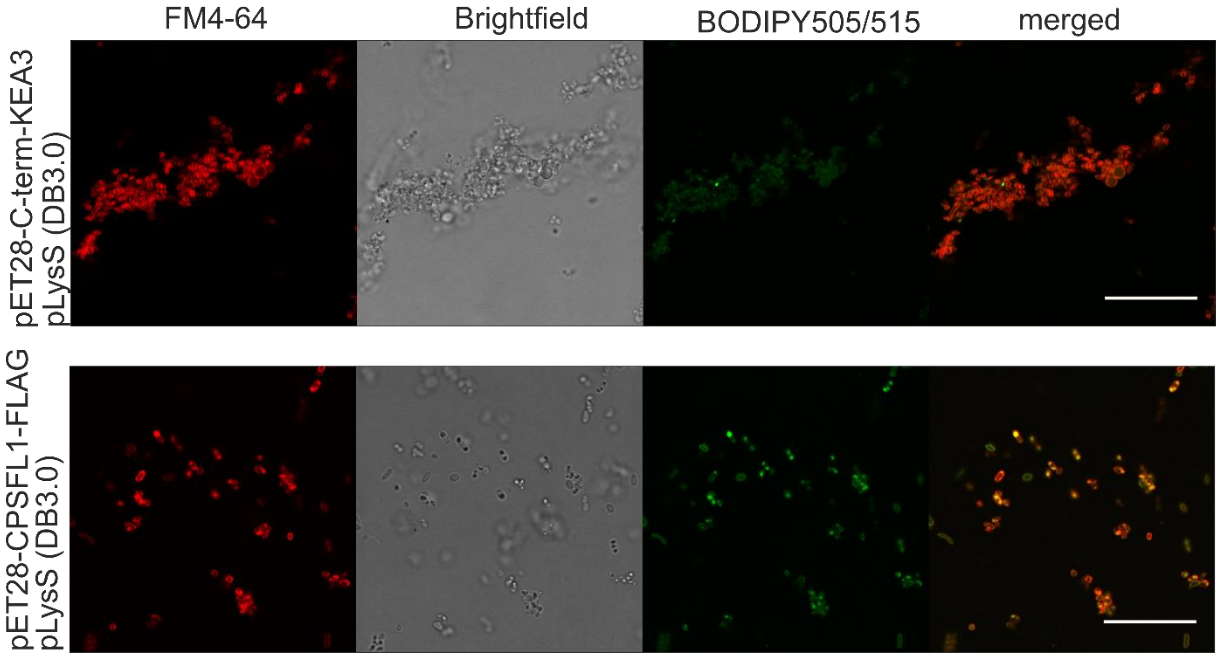


**Supplementary Figure 4**

Overview image of FM4-64 and BODIPY stained *E. coli* cells expressing a control protein (KEA3 C-terminus) or CPSFL1.

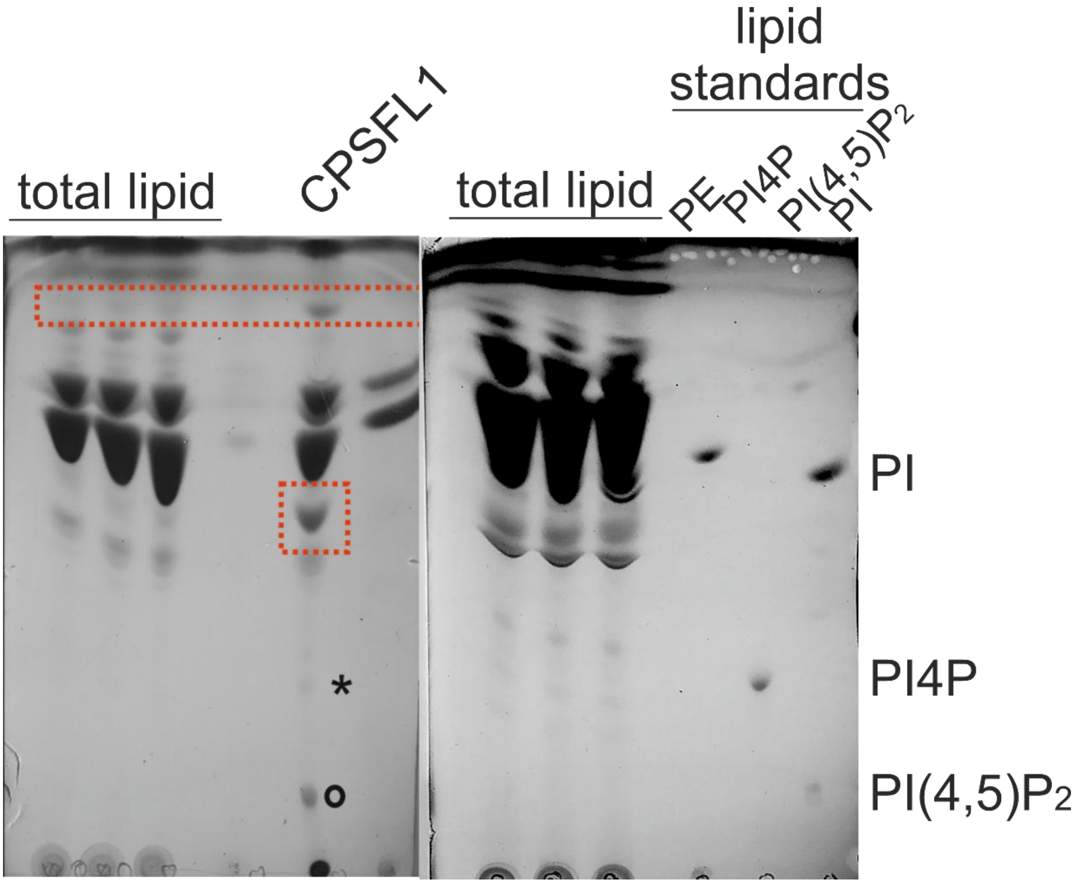


**Supplementary Figure 5**

TLC of PIP expressing cells and native extracts of CPSFL1 following Eosin Y endocytosis. Comparison with lipid standards identified PI4P and PI4,5P2 in the native CPSFL1 fractions. Overlay with the Eosin position indicated a prominent band (red square) enriched in native extracts as an Eosin derivate (right panel).

a

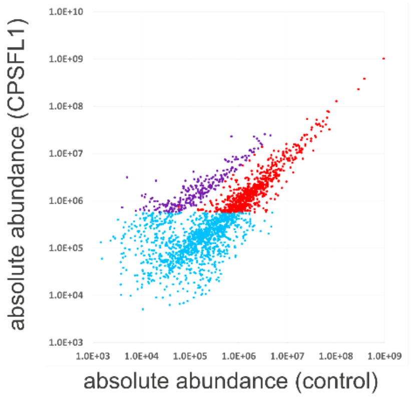


| | [n] | all features [n] | [%] |
|---|---|---|---|
| anotations in total | 850 | 3296 | 25.8 |
| anotations of abundant features | 280 | 3296 | 8.5 |
| anotations of interesting features | 130 | 3296 | 3.9 |
| | [n] | abundant features [n] | [%] |
| anotations of abundant features | 280 | 1000 | 28.0 |
| anotations of interesting features | 130 | 1000 | 13.0 |
| | [n] | interesting features [n] | |
| anotations of interesting features | 130 | 261 | 49.8 |

b

**Supplementary Figure S6**
This and the following.

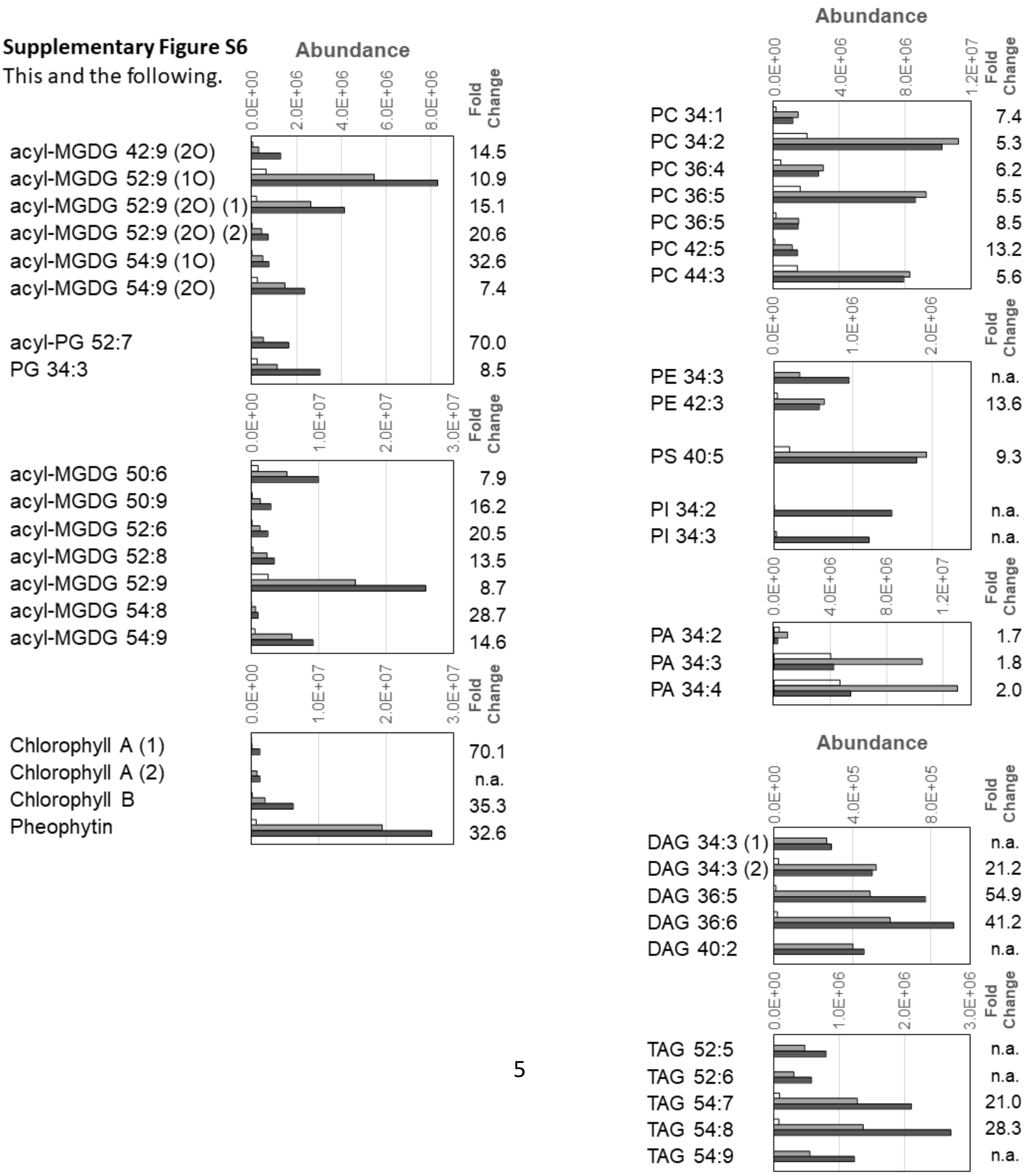

**Supplementary Figure 6**

**a,** Biplot of lipid mass features accumulated by co-immuno-purification together with YFP-tagged CPSFL1 from pre-purified chloroplasts of respective genetically modified plants relative a control lipid preparation of chloroplasts from plants that express YFP in chloroplasts. Non-targeted lipidomic analysis of the co-purified lipids revealed 261 mass features that accumulated at least 5-fold and were among the top 1500 abundant (arbitrary units) of 3294 detected mass features (violet), The abundance of most mass features remained unchanged (red) relative to the control. Low abundant mass features were omitted from further analyses (blue).

**b,** Characterization of the lipid fraction that was co-immuno-purified together with YFP-tagged CPSFL1 from pre-purified chloroplasts of respective genetically modified plants, Fig. 6c continued (refer to legend of Figure 6c). Acyl-MGDGs and Acyl-PGs include putative acylated lipids and Arabidopsides that were annotated according to expected monoisotopic exact masses; these compounds were not further characterized due to lack of reference substances. Diacylglycerol (DAG), triacylglycerols (TAG), phosphatidic acids (PA), phosphatidylinositol (PI), phosphatidylethanolamine (PE), and phosphatidylserine (PS). Note that we included PE and PS as controls of potential residual contaminations of the chloroplast preparation by eukaryote membrane lipids.
